# Pathogenic *Bacteroides fragilis* strains can emerge from gut-resident commensals

**DOI:** 10.1101/2024.06.19.599758

**Authors:** Renee E. Oles, Marvic Carrillo Terrazas, Luke R. Loomis, Maxwell J. Neal, Mousumi Paulchakrabarti, Simone Zuffa, Chia-Yun Hsu, Adriana Vasquez Ayala, Michael H. Lee, Caitlin Tribelhorn, Pedro Belda-Ferre, MacKenzie Bryant, Jasmine Zemlin, Jocelyn Young, Parambir Dulai, William J. Sandborn, Mamata Sivagnanam, Manuela Raffatellu, David Pride, Pieter C. Dorrestein, Karsten Zengler, Biswa Choudhury, Rob Knight, Hiutung Chu

## Abstract

*Bacteroides fragilis* is a prominent member of the human gut microbiota, playing crucial roles in maintaining gut homeostasis and host health. Although it primarily functions as a beneficial commensal, *B. fragilis* can become pathogenic. To determine the genetic basis of its duality, we conducted a comparative genomic analysis of 813 *B. fragilis* strains, representing both commensal and pathogenic origins. Our findings reveal that pathogenic strains emerge across diverse phylogenetic lineages, due in part to rapid gene exchange and the adaptability of the accessory genome. We identified 16 phylogenetic groups, differentiated by genes associated with capsule composition, interspecies competition, and host interactions. A microbial genome-wide association study identified 44 genes linked to extra-intestinal survival and pathogenicity. These findings reveal how genomic diversity within commensal species can lead to the emergence of pathogenic traits, broadening our understanding of microbial evolution in the gut.

## INTRODUCTION

The human microbiota is a complex community of microorganisms critical for host health. As one of the most abundant species in the human gut, *Bacteroides fragilis* is integral to various physiological processes including the regulation of metabolic processes and immune development. Colonization of the gastrointestinal tract by *B. fragilis* occurs shortly after birth and is detectable within the first few days of life^1,2^, with a single dominant strain establishing residence^3,4^. Although *B. fragilis* generally functions as a beneficial commensal throughout life, it can become pathogenic and cause opportunistic infections if it breaches the intestinal barrier. In fact, *B. fragilis* is the most common cause of anaerobic infections in humans^5–7^. Given the dual nature of *B. fragilis*^8^, we sought to elucidate the genomic features that distinguish commensal strains in the gut from pathogenic strains found in extra-intestinal environments.

Pioneering studies have explored the duality of commensal and pathogenic *B. fragilis* strains^6,9^. Despite extensive efforts, conclusive associations between clinical strains and specific virulence factors have yet to be established^10–14^. Some reports suggest that the capsule mediates serum resistance, agglutination, or adherence properties in virulent *B. fragilis* strains isolated from abscesses and bloodstream infections^11–13,15,16^. Purified capsular material from *B. fragilis* is sufficient to induce abscess formation in animal models^9,17^, highlighting its potential as a major virulence factor. On the other hand, capsular polysaccharide A (PSA) is known to govern immune tolerance in the gut^18^. Additionally, a subset of intestinal *B. fragilis* known as enterotoxigenic *B. fragilis* (ETBF), which produce an endotoxin encoded by the *bft* gene, are associated with infantile diarrhea and colorectal cancer^19–21^. The presence of these factors underscores the complex nature of *B. fragilis* as a key commensal in the gut microbiota that can turn pathogenic, leveraging its genetic and structural components to adapt and thrive in extra-intestinal environments. *B. fragilis* may adapt to challenging environments by evolving altered colonization capacities, novel metabolic capabilities, and interactions with different microbial communities to ensure survival. This duality poses significant challenges in understanding the precise mechanisms through which commensal strains of *B. fragilis* become pathogenic.

Here, we performed whole-genome sequencing for a comprehensive genomic and functional analysis of 813 commensal and pathogenic *B. fragilis* strains. This analysis unveils a vast and dynamic pangenome, illustrating the species’ remarkable adaptability to diverse host-associated niches and its transition from a commensal organism to a pathogen. We hypothesize that commensal strains act as reservoirs for virulence determinants, playing a crucial role in the emergence of pathogenic extra-intestinal isolates. Understanding this transition is critical for elucidating the pathogenic mechanisms associated with *B. fragilis* infections and underscores the need for further research into this complex commensal bacterium.

## RESULTS

### Comparative genomic analysis of commensal and extra-intestinal *B. fragilis* strains reveals extensive genetic diversity

We conducted the largest comparative genomic analysis to date of 813 *B. fragilis* whole genome sequences, which includes 147 newly isolated and/or sequenced strains (**Table 1**) and 666 sequences from public repositories ^22^ (**Table 2**). In total, 510 strains were derived from the gastrointestinal tract, 221 originated from extra-intestinal infections, and 82 from unknown origins. These strains were collected between 1925 and 2022, originating from 29 countries across six continents, and isolated from various anatomical sites including feces, intestinal biopsies, blood, and skin. We constructed a phylogenetic tree of all 813 isolates from the alignment of 2,349 orthologous core genes (found in ≥99% of *B. fragilis* strains), annotated with geographical and anatomical origins, and the presence and absence of select genes of interest (e.g., type VI secretion system (T6SS) GA1 and GA3, *B. fragilis* toxin) and plasmids (**Figure 1A**). Strain relatedness through the phylogenetic tree did not significantly correlate with either the continent of isolation (PERMANOVA, p=0.087), nor the site of isolation (intestinal vs. extra-intestinal; PERMANOVA, p=0.133), which corroborates previous studies^23^. This supports the hypothesis that pathogenic *B. fragilis* strains can emerge from gut-resident commensals through multiple independent events^24–26^.

**Figure 1.**
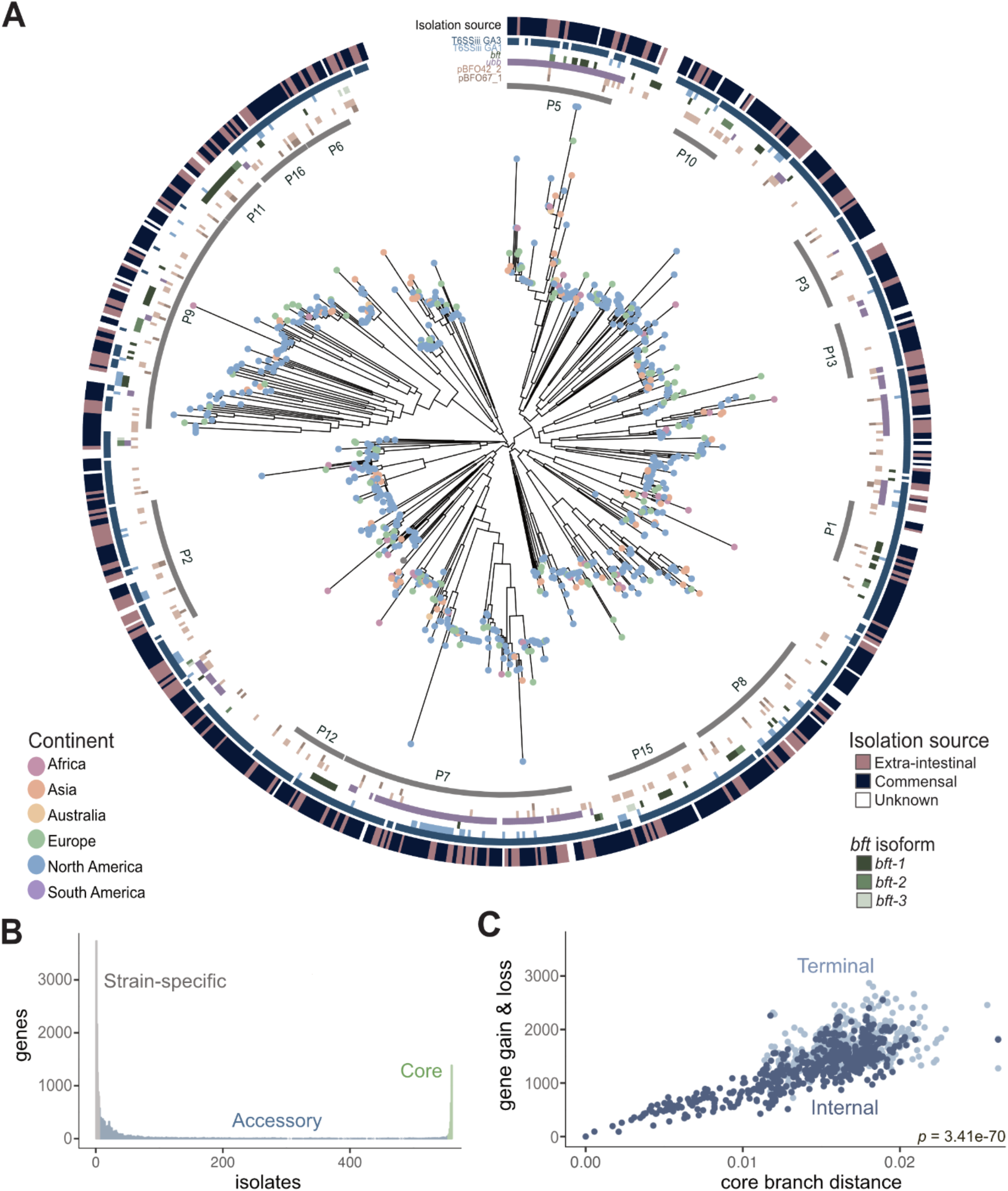
Pangenomic analysis of *Bacteroides fragilis* strains reveals extensive genetic variation. **A)** A core genome alignment-derived phylogenetic tree of *B. fragilis* strains by maximum-likelihood annotated with continent of isolate origin (located as dots at the tips), specific phylogroup clusters indicated by the gray inner ring, genetic regions of interest (the six colored rings which signify Type VI secretion system (T6SS) GA1 and GA3, pBFO42_2 and pBFO67_1, plasmid; *ubb; bft, B. fragilis* toxin with the three isoforms as separate shades of green (n=813; p-value of PERMANOVA of anatomic site = 0.133; p-value of PERMANOVA of geography = 0.087) with the anatomic site of isolation coloring the outermost ring. **B)** The number of genes that are strain-specific (in less than 1% of isolates), accessory (between 1% and 99% of isolates), or core (in more than 99% of isolates) in the *B. fragilis* pangenome of 813 strains. **C)** A measure of core compared with accessory genome evolution through cumulative gene gain/loss events (accessory genome) versus cumulative phylogenetic branch distance (core genome) (Student’s t-test, p = 3.41e-70), colored by terminal (light blue) and internal branches (dark blue) using the Panstripe algorithm (Tonkin-Hill et al., 2023).

In order to characterize the composition of the pangenome, we analyzed the genes of all 813 *B. fragilis* isolates and categorized them into core genes (2,349 orthologous genes found in ≥99% of *B. fragilis* strains), accessory genes (10,724 orthologous genes in <99% and >1%), and strain-specific genes (12,121 genes present in ≤1% of strains) (**Figure 1B and Table 3**). On average, a given *B. fragilis* isolate harbors 4,496 genes (± 273, standard deviation), composed of 52% core and 48% accessory genes. Core genes are found predominantly in conserved essential pathways, whereas accessory genes are commonly found in pathways facilitating niche-specific functions such as those involved in metabolic capabilities or immune evasion^27^ (**Figure S1A-1B**). We observed no significant differences in the number of core and accessory genes between gut-associated *B. fragilis* and extra-intestinal strains (**Figure S1A-B**), nor differences in genome size or GC content (**Figure S1C-1D**). We next evaluated the rate of gene gain and loss in relation to core gene evolution, finding a significant correlation between core genome phylogeny branch length and gene gain and loss events (Student’s t-test, p=3.4e-70) (**Figure 1C**). This indicates that *B. fragilis* exhibits considerable genetic diversity and proficiently assimilates new genes, a characteristic often observed in bacteria within diverse communities^28^. The extensive accessory genome of *B. fragilis* highlights its genomic plasticity, which is crucial for the acquisition of new genetic determinants that support the transition from a commensal lifestyle to survival in extra-intestinal niches^29^.

### *B. fragilis* phylogroups harbor distinct genes that mediate capsule composition and niche occupancy

Bacterial species are categorized based on shared evolutionary lineages, often reflecting their specific ecological niches^30–33^. In the case of *B. fragilis,* we employed a k-mer clustering approach using full genome assemblies to classify strains into phylogroups, revealing 16 distinct phylogroups (**Figure 2A, S2A, and Table 4**). Extra-intestinal *B. fragilis* strains are distributed across all phylogroups (**Figure S2B**), supporting the notion that pathogenic strains emerge independently from each phylogroup. Notably, we observed that commensal strains dominate phylogroups 10 and 15, while extra-intestinal strains are enriched in phylogroups 2 and 11 (**Figure S2B**).

**Figure 2.**
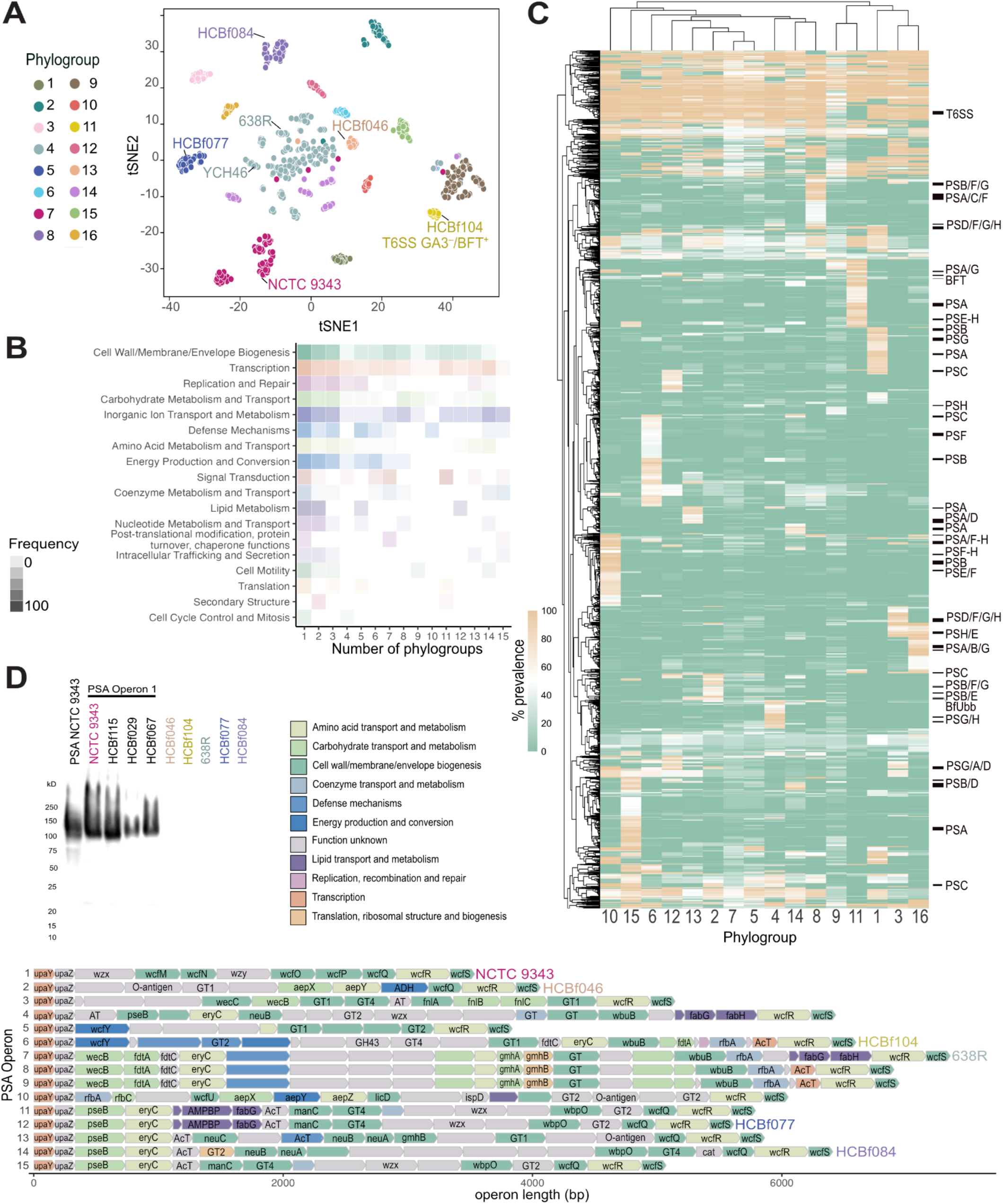
*B. fragilis* phylogroups have unique core-phylogroup genes. **A)** *B. fragilis* phylogroups based on t-distributed Stochastic Neighbor Embedding (t-SNE) algorithm of a MASH, k-mer based distance matrix of the whole genome sequence of all strains. Phylogroups labeled 1-16. The commonly used laboratory strains, NCTC 9343, YCH46, and 638R, are labeled within their respective phylogroups along with other *B. fragilis* strains used in functional assays described in this study: HCBf046, HCBf084, HCBf077, and HCBf104. T6SS GA3^−^/BFT^+^ represents a cluster of strains that lack T6SS GA3 and are positive for the presence of *bft*. **B)** Proportion of genes that are core to a collection of phylogroups, between 1 and 15, divided into COG categories with proportion expressed by the alpha of the heatmap cells. **C)** A heatmap of gene prevalence in each phylogroup. A euclidean clustering algorithm was applied to the rows and columns, gene clusters that belong to capsular polysaccharide paths are labeled, as well as the T6SS GA3 gene cluster and the location of *bft* and BfUbb. **D)** Western blot analysis of PSA of *B. fragilis* strains representing six PSA operon structures. Rabbit anti-PSA antibody was raised against NCTC 9343, which representing PSA operon 1 (top left). PSA operon structure of *B. fragilis* strains of high-quality assemblies (n=262). Genes colored by COG category and annotated with a gene name, if available by Bakta annotation.

Differential gene presence/absence analysis across phylogroups identified distinct gene clusters specific to certain phylogroups. The predominant category of core phylogroup genes is associated with the biosynthesis of cell wall and membrane, with a substantial proportion of genes related to capsular polysaccharides (**Figure 2B-2C**). *B. fragilis* capsular polysaccharides, particularly polysaccharide A (PSA), play a pivotal role in immune tolerance within the gut^18,34,35^, yet it also acts as a virulence factor involved in abscess formation outside the gut^9,17,36^. Previous studies have revealed structural variation in PSA among lab strains^37–39^. In our analysis of 262 high-quality genome assemblies (**Table 5**), we discovered 15 configurations of the PSA operon, characterized by conserved start (*upaY*, *upaZ*) and end (*wcfS*) genes^17,38^, but with distinct variation in intermediary genes (**Figure 2D**). These variations, designated operons 1-15, closely align with the 16 identified phylogroups (**Figure 2D, S2C**). A similar pattern is seen in the other capsular polysaccharides (PSB-PSH) (**Figure S3**). We observed that PSA operon 2 is enriched with commensal *B. fragilis* strains, whereas PSA operons 7 and 9 are dominated by extra-intestinal strains (**Figure S2D)**. Notably, the well-studied PSA structure from NCTC 9343 (referred to as PSA operon 1) is present in only 14.8% of the strains we examined (**Figure 2D, S2C**). Given its relevance in promoting immune tolerance, we sought to explore the structural similarities among PSA operons derived from representative *B. fragilis* strains by elucidating their cross-reactivity with the previously described PSA operon 1 from NCTC 9343. Western blot analysis indicated that antibodies raised against NCTC 9343 PSA can detect other *B. fragilis* strains harboring the PSA operon 1 (**Figure 2D, top left**). However, no cross-reactivity was detected to strains from five different PSA operons, indicating distinct antigenic variation among them. Overall, the genetic diversity within PSA operons contributes to antigenic variability, which may influence the balance between the induction of intestinal immune tolerance and extra-intestinal abscess formation under various contexts and niches.

Transmission electron microscopy analysis of *B. fragilis* strains also revealed morphology and ultrastructural variation across different strains and phylogroups (**Figure 3A**). This phenotype is further supported by monosaccharide and fatty acid analysis of capsular polysaccharide and lipopolysaccharide (LPS), revealing quantitative variation in monosaccharide composition among different *B. fragilis* strains (**Figure 3B-C and Table 6**). We did not observe repeating oligosaccharide chains typical of LPS, as previously reported^40,41^. Instead, we detected minor variations of lipooligosaccharide (LOS) structures in *B. fragilis* strains over six different phylogroups (**Figure 3D**). We further reveal an enrichment of *rfb* genes, involved in O-antigen synthesis, in extra-intestinal strains (**Figure S2E**). The modification of O-antigen expression in extra-intestinal strains, known to impact bacterial immune evasion through processes such as serum/complement mediated killing and niche occupancy^42^. To assess the potential impact of O-antigen variation in host response, we examined resistance to killing by normal human serum (**Figure 3E**). The extra-intestinal strain HCBf084 isolated from a wound drainage was highly resistant to serum killing. Notably, the survival of type strain NCTC 9343 was decreased in PBS alone, thus its susceptibility to human serum could not be evaluated under the same conditions. Altogether, these findings support that structural variations can impact the pathogenicity and virulence traits of *B. fragilis* strains.

**Figure 3.**
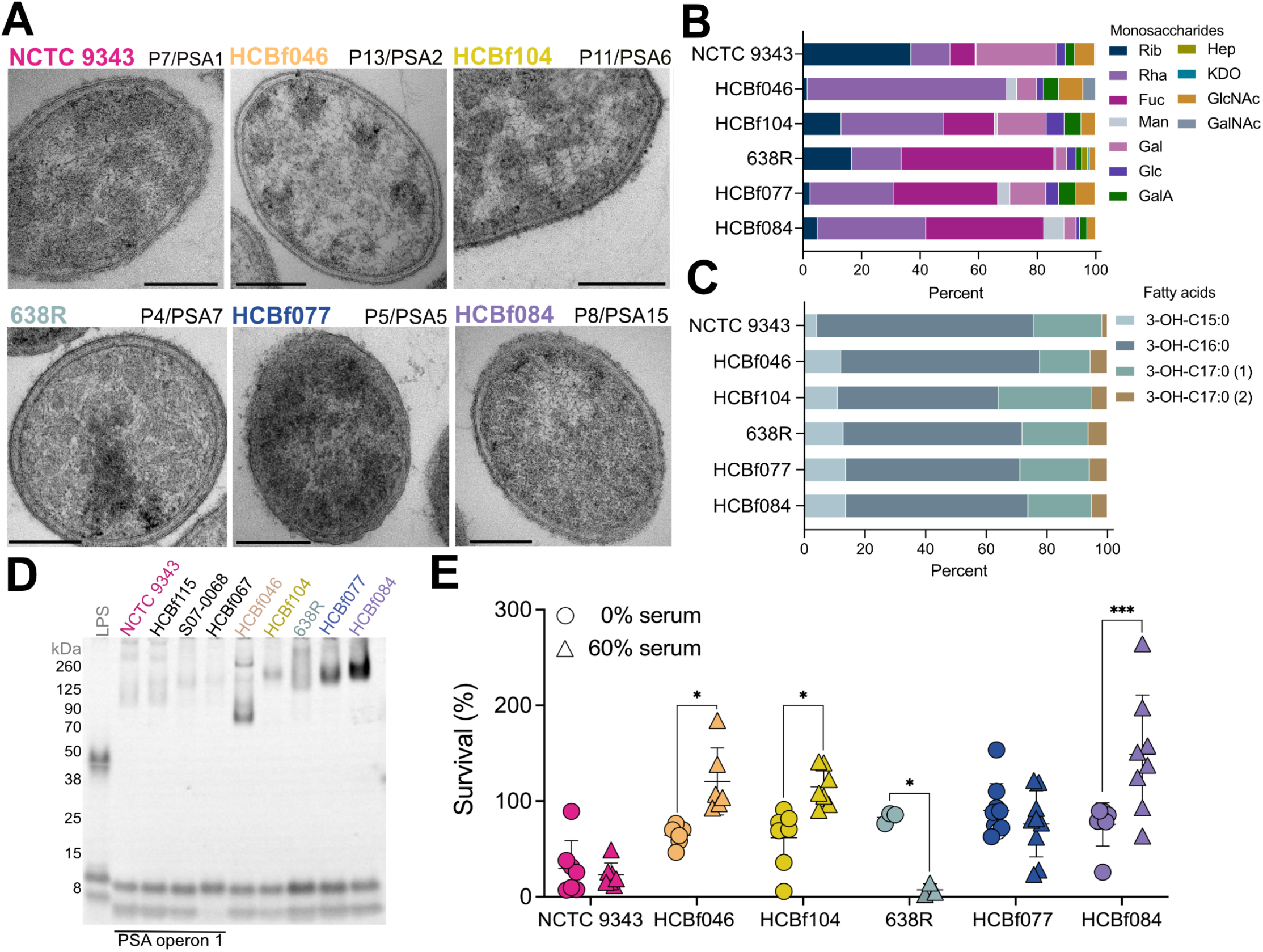
Capsule heterogeneity among *B. fragilis* strains contributes to variable host immune responses. **A)** Transmission electron microscopy of six representative *B. fragilis* strains: NCTC 9343, HCBf046, HCBf104, 638R, HCBf077, and HCBf084. Phylogroup and PSA operon (P#/PSA#) are as indicated. Scale bar, 200 nm. **B)** Capsule analysis where mole percentage of monosaccharides of six representative *B. fragilis* strains was determined by GC-MS. Rib, ribose; Rha, rhamnose; Fuc, fucose; Man, mannose; Gal, galactose; Glc, glucose; GalA, galacturonic acid; Hep, heptose; KDO, ketodeoxyoctonic acid; GlcNAc, N-acetylglucosamine; GalNAc, N-acetylgalactosamine. **C)** Capsule analysis where fatty acid percentage of six representative *B. fragilis* strains was determined by GC-MS. **D)** O-antigen analysis of whole cell lysates from six representative *B. fragilis* strains stained using Pro-Q Emerald glycoprotein stain. **E)** Serum killing assay of six representative *B. fragilis* strains. 10^6^ CFU of *B. fragilis* were treated with 0% or 60% normal human serum (NHS) in 1x PBS for 180 minutes. Results show the percent survival of *B. fragilis* strains. Two-way ANOVA comparing the mean of *B. fragilis* strains survival at 0% and 60% NHS. Data are representative of 3 experiments.

Beyond the capsule-related genes, our analysis of differential gene presence/absence between phylogroups revealed distinct gene clusters present in specific phylogroups (**Figure 2C and S2F**). Phylogroup 11 is enriched with extra-intestinal strains, representing 44% of this phylogroup (relative to 30% of total extra-intestinal strains in this study) (**Figure S2B**), with all the strains harboring the *B. fragilis* toxin (*bft*) gene (**Figure 1A and 2C**). We also observed that the T6SS GA3 (BF9343_1919-1925, 1931, 1940-1943) is absent in phylogroup 11 (**Figure 1A and 2C**). The *B. fragilis* T6SS GA3 system is known for its role in colonization and competition^43,44^. Therefore, its absence in all of phylogroup 11 is notable. The consistent presence of *bft+* strains within phylogroup 11 suggests a potential compensatory role in colonization and long-term niche occupancy, as previously hypothesized^45,46^. Additionally, the T6SS loci of GA3 is highly divergent, with distinct effector region variants aligning with phylogroups (**Figure S4**), and mirrors the pattern observed for the capsular polysaccharide paths (**Figure 2D and S3**)^47–49^. *B. fragilis* can also secrete a ubiquitin homolog (BfUbb) that targets select *B. fragilis* strains for lysis, mediating another mechanism of intra-species antagonism^50,51^. In our analysis, BfUbb (BF9343_3779) emerges as 11% more abundant in extra-intestinal strains and functions as a core gene within phylogroup 5. Moreover, we identified it in 87% of the strains belonging to phylogroup 7, while absent in other phylogroups (**Figure 1A and 2C**). Altogether, the distinct gene signatures identified within individual phylogroups indicate the potential for eliciting unique interactions between hosts and microbes.

### Growth and metabolic profiles of commensal and extra-intestinal *B. fragilis*

To determine if extra-intestinal strains exhibit distinct growth rates compared to commensal strains, we analyzed the growth of 78 phylogenetically representative *B. fragilis* strains in nutrient-rich media (BHI-S). Our analysis revealed no significant correlation between growth rate and isolation source (Welch’s t-test, p=0.6443), suggesting similar growth capabilities across commensal and extra-intestinal strains (**Figure 4A**). To further explore potential metabolic differences, we constructed genome-scale metabolic models for commensal and extra-intestinal strains. These models did not indicate any significant differences in metabolic potential between commensal and extra-intestinal *B. fragilis*, corroborating the growth assays (**Figure 4B**). Specifically, commensal and extra-intestinal strains could not be separated by predicted nutrient utilization profiles, predicted reaction rates to achieve optimal growth, or their individual metabolic enzyme and transporter inventories. Thus, neither the components of the models nor their predictions provide a basis for how commensal and extra-intestinal strains may be differentiated. Nonetheless, leveraging our whole genome data to construct these strain-specific models enabled accurate predictions of *B. fragilis* growth on diverse carbon and nitrogen sources (**Figure 4C**). This collection of models accurately predicts growth in 29 out of 33 conditions, which we verified through our growth assays or through public data^52^ on growth of *B. fragilis* strains (**Figure 4D**).

**Figure 4.**
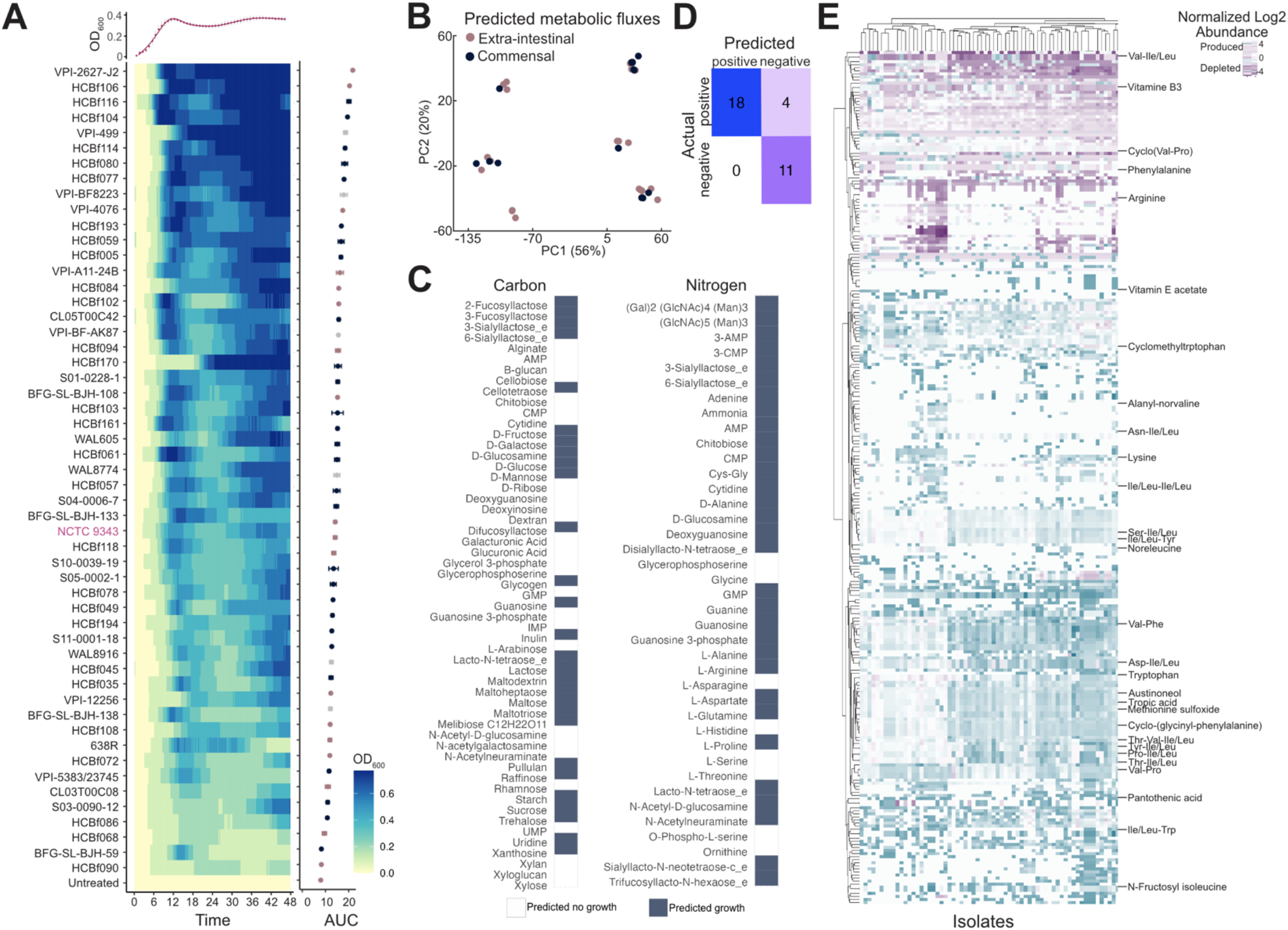
Growth and metabolic profiles of *B. fragilis* strains. **A)** Growth of *B. fragilis* strains over time and normalized to untreated wells, as indicated by OD_600_. Data is displayed as a heatmap, with a representative growth curve for NCTC 9343 overlaid above. Strains are ordered according to descending area under the curve (AUC). Data represents the AUC ± SEM, with the mean across three individual experiments. AUC coloring by extra-intestinal (pink), commensal (blue), and unknown (grey). **B)** PCA score plot of predicted metabolic fluxes, colored by commensal (blue) versus extra-intestinal (pink) strains. **C)** Defined carbon and nitrogen sources colored by growth (blue) or no growth (white) predicted for any *B. fragilis* strain. **D)** Predicted versus true growth of strains in defined nitrogen and carbon sources. **E)** Heatmap of differentially abundant metabolites compared to a media control, metabolites with annotation are labeled, hierarchical clustering of metabolites by row and strains by columns. Benjamini-Hochberg adjusted p ≤ 0.05 and log2-fold change ≥ 0.25; n of metabolites = 278; n of strains = 93.

We next performed untargeted metabolomic profiling of 84 *B. fragilis* strains (commensal n=38, extra-intestinal n=31, unknown n=18) via LC-MS/MS (**Figure 4E**). However, in line with the metabolic model, our analysis did not demonstrate clustering by isolation source (PERMANOVA, p=0.992) (**Figure S5A**). We did observe 12 metabolic features differentially produced between commensal and extra-intestinal strains (Wilcoxon rank-sum test, adjusted p≤0.05), however these hits were unannotated (**Figure S5B-C**). We also examined the prevalence of antimicrobial resistance genes (**Figure S6A-C**) and profiled the resistance activities in commensal and extra-intestinal isolates (**Figure S6D-F and Table 7**). Again, no significant differences in antimicrobial resistance were observed between commensal and extra-intestinal strains, demonstrating that resident intestinal strains may serve as a reservoir for antimicrobial resistance genes for the emergence of pathogenic strains outside the gut environment.

### A genome-wide association study identifies extra-intestinal-associated traits

To further understand the genetic factors contributing to pathogenicity and adaptation to extra-intestinal environments, we next profiled the accessory genome to identify genes associated with the extra-intestinal niche. We performed a microbial genome-wide association study (mGWAS) (n=514 non-MAG isolates) and uncovered 44 genes associated with isolation source (Holm’s-corrected p-value ≤ 0.05) (**Figure 5A**). Notably, our mGWAS analysis identified more gene associations with extra-intestinal isolates (Groups 2-8) than commensal strains (Group 1) (**Figure 5B and Table 8**). The mGWAS analysis identified a gene cluster associated with commensal strains (Group 1, light green; **Figure 5B**), with the majority of the genes linked to conjugative transposons CTn86 and CTn9343^53^. This region includes several conjugative transposon genes (e.g., *traK*, *traM*, *traN*), together with mobilization genes belonging to the BFT pathogenicity island (e.g., *bfmC, bfmA*). In *B. fragilis* toxin-expressing strains, the *bft* insertion site (BF9343_1444-1446) is located between the two mobilization genes significantly associated with commensal strains (**Figure 5B**, highlighted by pink borders)^54^. These findings align with the known associations of the *B. fragilis* toxin with gastrointestinal conditions such as diarrhea, inflammatory bowel diseases (IBD), and colorectal cancer^55–57^, supporting its prevalence in resident intestinal strains. Moreover, this region features a polysaccharide utilization loci (PUL) containing a SusC and SusD pair (BF9343_1437, BF9343_1436) and xylosidase (GH30; BF9343_1435). Adjacent to this PUL is a bile salt hydrolase (BF9343_1433) (**Figure 5B**), which a recent report linked bile acid metabolism to carbohydrate metabolism and PUL expression in *B. thetaiotaomicron*^58,59^. In our analysis, we identified three bile salt hydrolases in the *B. fragilis* pangenome, BF9343_3488 is core to the species, BF9343_1433 is in 47.23% (384/813), and HCBf132_07015 is present in only 0.023% (19/813) of strains.

**Figure 5.**
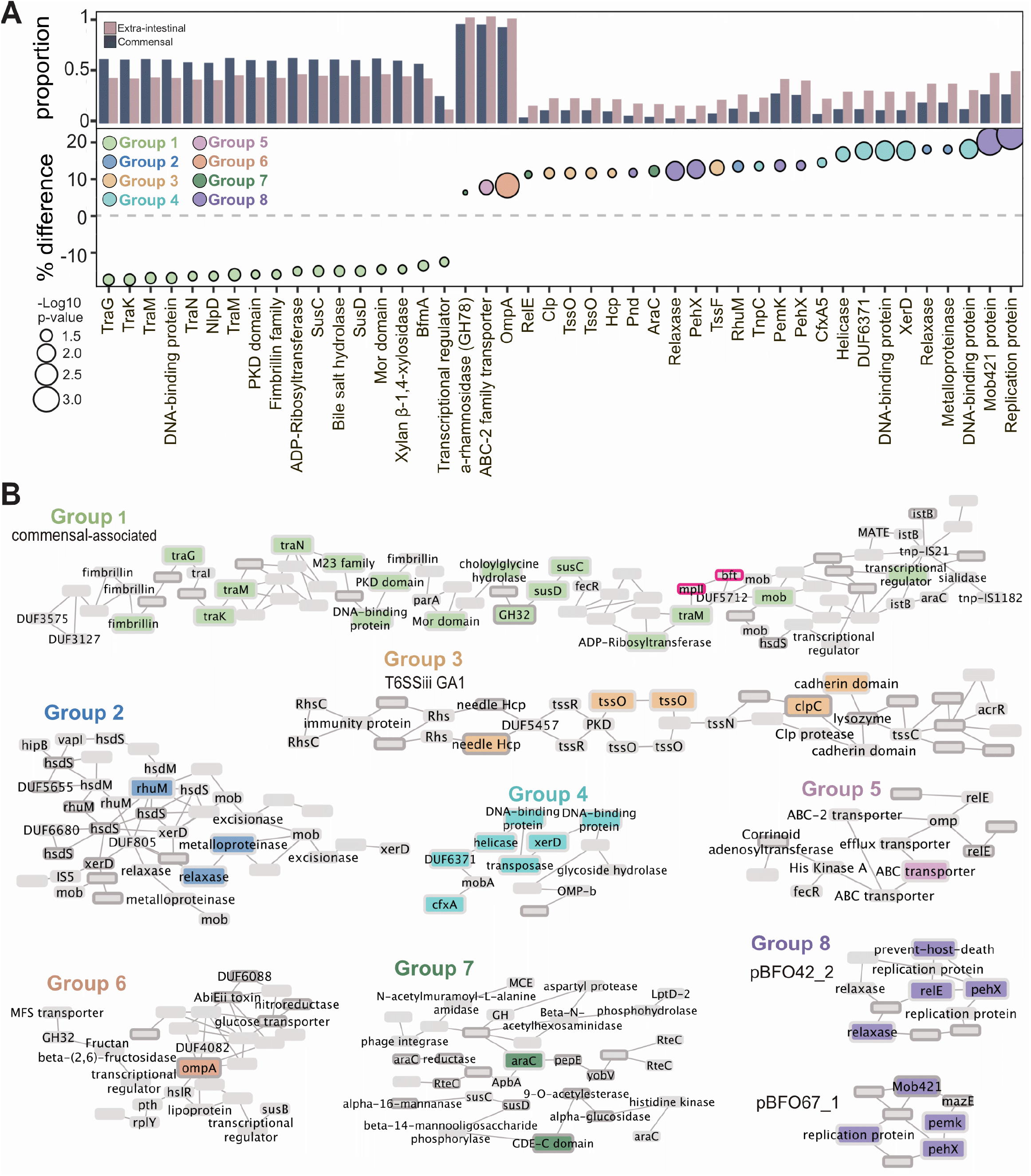
Extra-intestinal strains have genetic features associated with their isolation site. **A)** Genes identified through gene presence/absence mGWAS as differentially present in commensals versus extra-intestinal strains (dark blue, commensal; pink, extra-intestinal strains; Holm’s adjusted p≤0.05). Y axis is proportion of commensal or extra-intestinal strains with that gene as well as the percent difference in prevalence in commensal versus extra-intestinal strains. Size represents the -log10 p-value and color of each gene cluster determined by location on a pangenome graph, n=510 which excludes all metagenome assembled genomes (MAGs). **B)** Graphical representation of genes in the context of the pangenome in eight gene clusters. Group 1, commensal associated; Groups 2-7, extra-intestinal associated. The black border indicates genes which are predicted to be acquired through horizontal gene transfer (HGT). The pink border is highlighting the gene encoding the *B. fragilis* toxin, *bft*.

In extra-intestinal isolates, mGWAS identified several putative virulence genes (**Figure 5A-B**), including a metalloproteinase (BF9343_1074, Group 2). A higher abundance of genes associated with the T6SS GA1 system was also identified in extra-intestinal strains, particularly in the effector component, which includes Tss, Hsp, and Clp (Group 3, tan; **Figure 5B**). The T6SS GA1 locus, located on integrative conjugative elements, is widely distributed in gut *Bacteroidales*^47^ and suggested to be more readily transferred horizontally between species^60^. Strains harboring GA1 have a competitive advantage over other *Bacteroides*^61^. In extra-intestinal pathogenic *E. coli* (ExPEC), T6SSs play a role in virulence, conferring a highly pathogenic phenotype upon intestinal escape. Accordingly, extra-intestinal *B. fragilis* strains were associated with genes important for extra-intestinal survival (*araC*, Group 7) and response to stressors (*ompA*, Group 6), including antibiotics (*cfxA5*, Group 4; BF9343_2449, Group 5) (**Figure 5B**)^53,62^. Altogether, these genes enriched in extra-intestinal strains contribute to *B. fragilis* resistance to host immunity, facilitating successful intestinal escape and pathogenicity.

We also identified a set of accessory genes encoded on plasmids predominantly found in extra-intestinal *B. fragilis* strains (Group 8, purple; **Figure 5A-B**). The plasmid pBFO42_2^63^ was found in 40% of extra-intestinal strains, compared to 27% of commensals, while pBFO67_1^63^ occurred in 18% of extra-intestinal strains and only 5% of commensal strains. Both plasmids have been identified in other *Bacteroidaceae* including *B. uniformis, B. faecalis,* and *Phocaeicola vulgatus,* indicating their potential for interspecies transfer. Additionally, the broad distribution of plasmid pBFO42_2 across the phylogenetic tree suggests acquisition via horizontal gene transfer (HGT) (**Figure 1A**). These plasmids both harbor a toxin-antitoxin system and a putative exo-poly-alpha-D-galacturonosidase (GH28), an enzyme responsible for hydrolyzing alpha-D-galactose residues (**Figure 5A**)^64,65^. Consistently, we observed elements linked to extra-intestinal environments, either identified or located near mobilization genes and genes predicted to be acquired through HGT. Specifically, groups 1, 3, 5, and 8 contain at least one transposon or plasmid-associated gene (**Figure 5B**). Given that HGT is the primary mechanism for gene acquisition in prokaryotes^66^, these results suggest that *B. fragilis* strains have acquired genes horizontally, enhancing their fitness and adaptability to colonize new ecological niches, such as extra-intestinal environments^67–69^.

## DISCUSSION

This comprehensive genomic analysis of 813 *B. fragilis* strains reveals new insights into the genomic underpinnings that facilitate the commensal-to-pathogen transition. Our findings demonstrate an expansive pangenome, characterized by extensive genetic diversity that supports both the adaptability and the pathogenic potential of *B. fragilis*. Specifically, the identification of distinct phylogenetic groups associated with various virulence factors, including the *B. fragilis* capsule, T6SS, and toxin, offers new perspectives on how environmental pressures and HGT contribute to the evolutionary trajectory of these gut bacteria. Interestingly, many of these so-called ‘virulence factors’ also play crucial roles in immune regulation^18,70^ and colonization in the gut^60,71^. This dichotomy suggests that these factors may be more appropriately considered as ’niche factors’—a concept proposed by Colin Hill^72^, which are essential for survival and adaptation within specific environmental contexts rather than exclusively mediating pathogenicity. Our findings here support this notion, as phylogroup-specific genes were primarily associated with niche factors within *B. fragilis* strains (**Figure 2**). While historically associated with virulence, arguably the capsule, T6SS GA3, and even the BFT orchestrates some aspects of intestinal colonization and occupancy. This reconceptualization challenges the traditional pathogen-centric view of these genes and suggests a more complex interplay of bacterial survival strategies that include both symbiotic and pathogenic strategies.

Indeed, several studies have reported that enterotoxigenic *B. fragilis* (ETBF) strains evolved from non-toxigenic *B. fragilis* strains via horizontal transfer^21,73^. Phylogenetic analysis indicates that ETBF strains do not cluster together (**Figure 1A**) but likely emerged through multiple independent events. This, coupled with the open pangenome of *B. fragilis*, supports the notion that commensal strains act as reservoirs for virulence determinants, enabling the emergence of pathogenic strains once it escapes the normal gut environment. Our findings align with previous work on the commensal *E. coli* HS strain, which also exhibits genetic mosaicism with traits characteristic of both commensal and pathogenic strains^74^. Rasko and colleagues proposed that commensal bacteria may serve as ‘genetic sinks’ that give rise to pathogenic isolates. The dynamic nature of the *B. fragilis* pangenome, along with a higher rate of gene acquisition and HGT^69^, suggests that commensal strains continuously integrate new genetic elements, contributing to the emergence of pathogenic variants. These findings underscore the adaptability of *B. fragilis*, suggesting that factors often labeled as ’virulence factors’ are actually part of a broader adaptive strategy for navigating the complex gut environment, with implications ranging from commensalism to pathogenicity.

Our functional analyses further revealed indistinguishable growth capacities and metabolic profiles between commensal and extra-intestinal *B. fragilis*. However, when we profiled the accessory genome to identify associations with the intestinal versus extra-intestinal niche, we uncovered 44 genes linked to isolation sources using microbial genome-wide association study (mGWAS). This analysis identified more gene associations with extra-intestinal isolates than commensal strains, predominantly characterized as putative virulence genes essential for survival outside the gut. Notably, genes associated with commensal strains were primarily acquired via conjugative transposons, including CTn86 and CTn9343 (**Figure 5A**), which are found almost exclusively in *B. fragilis* division I strains^75,76^. These genetic elements associated with commensal *B. fragilis* have a very narrow host range relative to those found in extra-intestinal strains. In contrast, genes enriched in extra-intestinal isolates were acquired horizontally from strains within and beyond *Bacteroides*, including plasmids pBFO42_2 and pBFO67_1 (**Figure 5A-B**). A recent study reported the prevalence of the cryptic plasmid pBI143 in gut microbiomes from humans living an industrialized lifestyle, characterized by genes responsible for mobilization (*mobA*) and replication (*repA*)^77^. Although pBI143, initially described in *B. fragilis*^78^, does not appear to confer direct beneficial functions, its copy numbers increased during stress conditions such as oxygen exposure and IBD. This pattern is consistent with our findings on the prevalence of pBFO42_2 and pBFO67_1 in extra-intestinal *B. fragilis* isolates, suggesting that these plasmids may provide adaptive advantages under environmental stresses typical of extra-intestinal infections, such as increased oxygen exposure.

Altogether, this study elucidates the adaptive mechanisms enabling *B. fragilis* to transition between commensal and pathogenic roles, highlighting the intricate interplay of genetic, environmental, and evolutionary forces shaping this duality.

## LIMITATIONS

While the large scale of our genomic dataset provides robust statistical power, the functional roles of many identified genes remain to be elucidated through experimental validation. Future research should focus on longitudinal studies to track the evolutionary dynamics of *B. fragilis in situ*, examining how shifts in the gut environment—such as changes in diet, antibiotic usage, or disease states—affect its genomic architecture and pathogenic status. Additionally, exploring the interaction networks between *B. fragilis* and other gut microbiota members could shed light on the community-level mechanisms that favor its pathogenic transformation.

## ACKNOWLEDGEMENTS

We thank members of the Chu lab for technical support and helpful discussions. We also thank A. Russell, G. Sharon, and A. Khosravi for valuable feedback and discussions. This work was supported by grants from the National Institute of Health (NIH) R01 AI167860 and P30 DK120515. Support for R.E.O. was provided by T32 AR064194 (NIAMS) and M.C.T by T32 DK007202 (NIDDK), the National Academies of Sciences, Engineering and Medicine through the Predoctoral Fellowship of the Ford Foundation, and the Howard Hughes Medical Institute (HHMI) Graduate Fellowships grant (GT15123). M.H.L. was supported by T32 DK007202 and F32 AI169989. M.R. was funded by the NIH grants AI126277, AI145325, and the UCSD Department of Pediatrics. Additional support was provided to M.R. and H.C. by the Chiba University-UC San Diego Center for Mucosal Immunology, Allergy and Vaccines (cMAV) and AMED (JP233fa627003). The authors thank the University of California, San Diego - Cellular and Molecular Medicine Electron Microscopy Core (UCSD-CMM-EM Core, RRID: SCR_022039) for equipment access and technical assistance. The UCSD-CMM-EM Core is partly supported by the National Institutes of Health Award number S10OD023527. This publication includes data generated at the UC San Diego IGM Genomics Center utilizing an Illumina NovaSeq 6000 that was purchased with funding from a National Institutes of Health SIG grant (#S10 OD026929).

## Conflict of Interest

W.J.S’s current conflicts of interest are: Mirador Therapeutics (stock, employee, company officer, Ventyx Biosciences (stock, former employee), Prometheus Laboratory (board of directors), Shoreline Biosciences (stock, scientific advisory board), Forbion (consultant), Alimentiv (consultant). P.C.D. is an advisor and holds equity in Cybele, BileOmix, and Sirenas and a Scientific co-founder, and advisor and holds equity to Ometa, Enveda, and Arome with prior approval by UC-San Diego. P.C.D. also consulted for DSM animal health in 2023. R.K.’s current conflicts of interest are: Gencirq (stock and SAB member), DayTwo (consultant and SAB member), Cybele (stock and consultant), Biomesense (stock, consultant, SAB member), Micronoma (stock, SAB member, co-founder), and Biota (stock, co-founder).

## METHODS

### Bacterial strains and culture conditions

Bacterial strains are described in Table 1. *Bacteroides fragilis* strain NCTC9343 was obtained from the American Type Culture Collection (ATCC). *Bacteroides fragilis* strains were grown anaerobically (10% H2, 10% CO2, 80% N2; Coy Lab Products) at 37 °C in brain heart infusion (BHI) broth (BD Biosciences) supplemented with 5 μg/ml hemin (Sigma) and 0.5 μg/ml vitamin K (Sigma) (BHI-S).

### Sample collection and bacterial isolation

Samples from healthy donors and patients were collected with the approval of the University of California San Diego Institutional Research Board and with written informed consent signed by subjects prior to sample collection. Healthy and IBD strains were collected under IRB #141853, #150675, and #190012, and extra-intestinal *B. fragilis* strains were collected from the UCSD Clinical Microbiology Facility under IRB# 160524. *B. fragilis* strains from healthy donors were generously provided by Dr. Eric J. Alm (Massachusetts Institute of Technology)^4^. The historical *B. fragilis* strains were curated from the laboratory of Dr. Abigail Salyers (University of Illinois at Urbana-Champaign)^25,79^ and provided by Dr. Eric Martens (University of Michigan)^52^. Public strains were obtained from the NCBI repository for *Bacteroides fragilis* samples. In addition, we analyzed samples from previously published studies^7,22^.

*B. fragilis* strains were isolated from approximately 0.5 g fecal materials, which were homogenized in 30% glycerol/0.1% cysteine, diluted 1:10, and plated in BHI-S with gentamicin (100 µg/mL). Colonies were picked from BHI-S plates and identities were determined using *Bacteroides* species-specific primers by qPCR (**Table 9**), followed by confirmation by Sanger sequencing using primers for 16S rRNA, 27F and 1492R^80^. *B. fragilis* strains were banked in glycerol and stored in -80 °C for downstream whole genome sequencing and functional studies.

### Metagenomic Library Preparation and Sequencing

*Bacteroides* strains were grown anaerobically for 72 hours at 37°C in 1 ml of BHI-S. Nucleic acid extraction was performed with the MagMAX CORE Nucleic Acid Purification Kit (ThermoFisher)^81^. Extracted genomic DNA was transferred from 2.0 mL Eppendorf tubes to 96-well plates, and then compressed into a 384-well plate. Using 1 µL of each sample, gDNA was quantified using the PicoGreen™dsDNA Assay Kit (ThermoFisher Scientific). Based on this quantification, each sample was normalized to 3.5 ng in 5 µL water before generating libraries using a 1:10 Kapa HyperPlus (Roche) miniaturized protocol with barcoded indices as previously described^82^. The libraries were quantified using the PicoGreen™ dsDNA Assay Kit (Thermo Fisher Scientific), and an equal volume of each sample was pooled using an Echo 550 acoustic liquid handler (Labcyte), PCR cleaned using the QIAquick PCR Purification Kit (QIAGEN), then size-selected to fragments of 300-700 bp using a PippinHT (Sage Science). The average fragment length was determined using a High Sensitivity D1000 Tapestation (Agilent), and this was used to calculate the average molarity of the pool. This pool was diluted to 90 pM and sequenced with paired-end 150 bp on an iSeq v2 (300 cycle) (Illumina). Raw reads generated from the iSeq were demultiplexed, and new normalized pooling values were calculated based on read counts^83^. These new normalized pooling volumes were pooled from the original libraries, PCR cleaned, and again size-selected to fragment sizes of 300 - 700 bp^83^. The iSeq-normalized pool was sequenced on an Illumina NovaSeq 6000 with 2 x 150 bp chemistry at the Institute for Genomic Medicine at UC San Diego.

### Post-Sequencing Data Processing

After raw sequence reads were generated from the NovaSeq, adapter trimming was performed by Fastp^84^. Human filtering was performed by Minimap2^85^ by alignment to one database containing human reference genome GRCh38 and PhiX, and a second database containing human reference genome CHM13 (ref 116,117). Resulting FASTQ were uploaded into Qiita^86^ study ID #14360 (https://qiita.ucsd.edu/public/?study_id=14360) or at EBI under accession number PRJEB76295 ERP160853. FASTQ files were further quality controlled via Fastp.

### Creation and analysis of the pangenome

Assembly, quality control, pangenomic analysis, and genome-wide associations studies were conducted with the package Panpiper^87^. Briefly, draft assemblies were assembled using Shovill v1.1.0^88^ (parameters --nocorr). Shovill filtered contigs with low coverage (<2x). Contigs were also filtered using BBMAP v38.93 if they were less than 500bp long. Quality control removed assemblies with less than 95% completeness, greater than 5% contamination by Checkm v1.1.3, less than 500 contigs, less than 95% of reads mapping to *B. fragilis*, and an average nucleotide identity to the reference 9343 *B. fragilis* strain less than 95% by FastANI v 1.32. Of the 953 *B. fragilis* strains that passed assembly quality filtering, 140 mapped at approximately 87% sequence identity to the *B. fragilis* type strain NCTC 9343^87^. Thus, these strains were excluded from further analysis in this study. We also removed strains which were isolated over multiple timepoints from the same host to decrease redundancy in our sample pool. This filtered the input to 813 samples.

Bakta v1.6.0^89^, AMRFinderPlus v3.11.14^90^, and EggnogMapper v2.1.11^91^, were used to annotate the high-quality assemblies. A pangenome from the annotated files was created using Panaroo v1.2.10^92^ (parameters --remove-invalid-genes --clean-mode strict -a core --core_threshold 0.98 --len_dif_percent 0.98 -f 0.7 --merge_paralogs - t 20 --refind_prop_match 0.5 --search_radius 5000). The pangenome size was estimated using the Infinitely Many Genes model as implemented in panaroo and the openness of the genome determined through gene gain and loss events compared with the branch lengths in the core genome using Panstripe, with significance determined through a Student’s t-test^93^. A phylogenetic tree based on the core genome alignment was created using FastTree2 v2.1.11^94^, IQTree2 v2.2.0.3^95^, and RAxML v8.2.12^96^. Using the distances from the midpoint rooted phylogeny created by RAxML v8.2.12, a distance matrix was created to account for lineage effects in the genome-wide association study. Mash v2.3^97^ was used to create a distance matrix between the samples using locality-sensitive hashing (parameters -s 10000). The resulting distance matrix was used to divide the strains into phylogroups, with a cutoff value of 0.44^98^. The distance matrix was transformed via tSNE, using the fit_transform method from skylearn in Python and colored by phylogroup. We identified all genes core to individual phylogroups defined by 95% or more presence in all isolates of a phylogroup. We plotted shared core genes between phylogroups by COG category using ggplot in R. We additionally plotted a heatmap of all genes and their average abundance in each phylogroup using pheatmap in R.

To conduct a bacterial genome-wide association study, three sources of genetic variation were used – unitigs, SNPs, and gene presence/absence patterns. These genetic variants were tested between phenotypes including fecal vs non-fecal source and antimicrobial resistance. In these, we excluded assemblies from MAGs in the gene presence/absence test as the MAGs have incomplete genomes and have lost any associated plasmids, leading to 514 isolate genomes. The association study was conducted using FastLMM as implemented in Pyseer v1.3.9^99^ using a kinship matrix derived from the phylogenetic tree from the core genome alignment to account for population structure. We filtered the resulting p-values by BH correction for the structural and gene presence/absence tests and filtered by a threshold calculated in Pyseer based on the number of patterns measured for the unitig test. We used a graph-based representation of the pangenome, where each node is a gene, and each edge represents two genes being adjacent in at least one isolate genome to determine the position of the significant genes in relation to each other in Cytoscape^100^. We selected four edges from each gene of interest which revealed all significant genes fall into 8 gene clusters within the pangenome; we filtered these gene clusters to only allow genes present in more than 5% of the population and edges present in more than 2.5% of the population.

We filtered the assemblies to the most high-quality assemblies (less than or equal to 10 contigs), creating a pangenome of just these assemblies (n=262) (Table 5). The pangenome output contains a graph object where each node is a gene, and each edge represents two genes being adjacent on an assembled contig. Each edge and node are annotated with the strains that have that gene or connection. Since the first and last gene in the polysaccharide biosynthesis operons are conserved, a graph traversal program can be used to find all the possible paths between the first and last gene. We created a python package Travis (https://github.com/rolesucsd/Travis.git) to traverse the pangenome graph from a start and stop gene and used this tool to characterize several polysaccharide biosynthesis operons (PSA-H) and annotate the genes composing the pathway to genetically determine the significance of variations.

### Western blot analysis of Polysaccharide A

100µl of overnight *B. fragilis* culture was spun down at 800 x g for 5 minutes. Bacterial pellets were resuspended in Laemmli Sample Buffer (Bio-Rad) and 1% 2-mercaptoethanol. Samples were incubated at 99 °C for 10 minutes. Electrophoresis was conducted with 4-20% Tris-Glycine Mini Protein Gel (Invitrogen) and Tris-SDS glycine running buffer at 120V for 2 hours using a Mini Gel Tank (Invitrogen). Transfer to the PVDF membrane was performed overnight at 10mA. Membrane was blocked 1 hour at room temperature (RT) in 5% non-fat dry milk in 1x PBS. Membrane was incubated with primary antibody (anti-PSA at 1:1000) for 1 hour at RT and washed three times with 0.05% PBS-Tween (Fisher Scientific). Incubation with secondary antibody (goat anti-rabbit IgG at 1:10,000, Millipore) was performed for 1 hour at RT, followed by three washes with 0.05% PBS-Tween. Reaction bands were detected using enhanced chemiluminescence (ECL) (Life Technologies) using Chemiluminescent Western Blot Imager Azure 300 (Azure Biosystems).

### Isolation and purification of LPS/LOS

LPS/LOS was isolated from bacteria using slightly modified hot phenol-water extraction method^101^. Briefly, bacterial cells were suspended in 20 ml ultra-pure water and sonicated for 3 min, placed in a 68 °C water bath and kept stirring for 10 min. Equal volume of pre-heated (68 °C) 90% aqueous phenol solution was added to the cell suspension. The mixture was stirred vigorously for 45 min at 68 °C and then cooled rapidly in an ice-water bath for 10min. The sample was centrifuged at 6000 RPM for 45 min at 10 °C. The phenol saturated aqueous layer from the top was collected in a 50 ml Falcon tube and the residual material was re-extracted once more with equal volume of aqueous layer collected from first extraction using pre-heated ultrapure water following the similar method as mentioned above. The phenol-saturated aqueous layer was dialyzed using 1,000 molecular-weight-cutoff regenerated cellulose dialysis tubing against DI-water for 3 days (with one change of 4L water per day). The samples were lyophilized and used for composition analysis by GC-MS.

### Monosaccharide and fatty acid analysis

Monosaccharide and fatty acid composition was analyzed using GCMS as TMS derivative. Briefly, known amounts of LPS/LOS samples were spiked with 1 µg of myo-inositol as internal standard and methanolized using 1M MeOH-HCl at 80 °C for 16h. The samples were then dried using nitrogen flush and N-acetylated using a mixture of methanol: pyridine: acetic anhydride at 100 °C for 1h. The reaction mixture was dried using nitrogen flush followed by TMS derivatization using Tri-Sil reagent (Thermo). The samples were extracted with hexane and injected on GCMS (Agilent Tech) equipped with Restek-5ms capillary column and ultra-pure Helium was used as carrier gas. The monosaccharides were quantified by comparing with known amounts of standards and fatty acids were presented as relative area percentages. Details of the instrumental method are as described (Leker et al. 2017).

### O-antigen assays

5×10^8^ CFU/ml of a *B. fragilis* (OD_600_=0.3) was washed twice in PBS and then resuspended in 200 µl of lysis buffer (2% 2-Mercaptoethanol, 2% SDS, 10% glycerol, and 0.1M Trizma HCl in water adjusted to pH 6.8). Samples were incubated at 95 °C for 10 minutes and then incubated with 40 µl of Proteinase K (Sigma) overnight at 55 °C. Lysates were prepared for electrophoresis with Laemmli Sample Buffer (Bio-Rad) and 7.5% 2-Mercaptoethanol. Electrophoresis was conducted using a Mini Gel Tank (Invitrogen), Novex Tris-Glycine Mini Protein Gel 4-20% (Invitrogen), and MES SDS Running Buffer (Invitrogen) at 25mA for 2 hours. O-antigen staining was then performed with Pro-Q Emerald 300 Lipopolysaccharide Gel Stain Kit (Invitrogen) following the manufacturer’s instructions. Gels were imaged with 302 nm UV transilluminator Workhorse Gel Imager Azure 200 (Azure Biosystems).

### Human serum killing assay

*B. fragilis* strains were cultured in BHI-S under anaerobic conditions at 37 °C overnight before being subcultured and normalized to 10^9^ CFU/mL by OD_600_ and washed twice with sterile PBS. Approximately 10^6^ CFU of each *B. fragilis* strain was treated with 60% normal human serum (Complement Technologies) for 180 minutes under anaerobic conditions and total CFU was enumerated by plating on BHI-S.

### *Bacteroides* growth assays

*Bacteroides fragilis* strains were grown anaerobically, a single colony was picked for each strain and inoculated in 5 ml of BHI-S broth and grown anaerobically for 18 hours. Overnight cultures were subcultured at 1:25 (v:v) in fresh BHI-S and grown to OD_600_ of 0.8, 5 μL of this culture was taken and inoculated into a 96-well plate with 200 μL of BHI-S media.

For defined carbon sources a minimal media was used, for which 5 mg/mL of the said carbon sources and 1.25 mg/mL of yeast extract were both added^102^. The media was then sterilized with a 22μm filter and 200μL added to a 96-well plate. Bacteria is similarly grown to OD_600_ of 0.8 once subcultured, then washed twice by spinning at 8000 rcf for 5 mins and resuspending in minimal media without any carbon sources. 5μL of the washed bacterial culture was used in inoculate before the plate was read (BioTek) at OD_600_ every 15 minutes for 48 hours. All data was finally processed using the gcplyr^103^ and growthcurver^104^ packages within Rstudio^105^.

### Genome-scale metabolic modeling

Genome-scale metabolic models for the strains of interest were constructed using *B. fragilis* model *i*MN674^106^ as a template. Such models are composed of the enzymes, transport proteins, and their associated reactions and metabolites within an organism’s genome. Thus, they represent the set of possible molecular transformations an organism may perform, thereby allowing for the prediction of what compounds may be made in what amounts from a simulated medium. In combination with a list of metabolites that make up the organism’s biomass, these models can be used to predict growth rates under different conditions.

The strain specific models were created using protein BLAST to identify which genes in the template model had orthologues in each strain’s genome, with a cutoff of 80% identity. BLAST was also used against all Gram negative bacterial models in the BiGG database^107^. The biomass reaction was unchanged from *i*MN674.

All simulations were performed in a medium of ammonia, hydrogen sulfide, heme, minerals, and inorganic phosphate, with unbounded uptake. In carbon source simulations, the source indicated was provided with a maximum uptake rate of 10 millimoles per gram dry weight per hour. Glucose and galactose uptake values were fitted to reflect the growth rates found experimentally. For nitrogen source simulations, glucose was provided at the fitted uptake value or the average of these values for strains without data. The nitrogen source was provided at 10 millimoles per gram dry weight per hour. All simulations were performed in MATLAB R2022b using the COBRA toolbox^108^ version 3.3.

### Extraction of bacterial cultures for LC-MS/MS

*Bacteroides* strains were grown anaerobically for 24 hours at 37 °C in 500μl of brain heart infusion broth (BD Biosciences) supplemented with 5 μg/ml hemin and 0.5 μg/ml vitamin K (BHI-S) and stored at -80 °C until subsequent analysis.

Samples underwent three freeze-thaw cycles to lyse the cells. A total of 180 μL were collected from each sample and transferred to a deep 96-well plate. Subsequently, 600 μL of LC-MS grade methanol (MeOH) was added to each well. The samples were then covered and sonicated for 10 minutes. After that, samples were centrifuged for 15 minutes at 2000 rpm. Supernatant (200 μL) was then transferred to a shallow 96-well plate to be evaporated in vacuum pressure under centrifugation. The dried samples were then stored at -80 °C until LC-MS/MS analysis. On the day of the LC-MS/MS experiment, dried extracts were resuspended in a 50% MeOH/H_2_O solution containing sulfadimethoxine (1 μM) as internal standard. Resuspended samples were then sonicated for 10 min before the analysis.

### LC-MS/MS analysis of culture extracts

Culture extracts (5 µL) were analyzed using a Vanquish ultra high-performance liquid chromatography (UHPLC) system coupled to a Q Exactive quadrupole orbitrap mass spectrometer (Thermo Fisher Scientific, Waltham, MA, USA). A Kinetex C18 column (50 x 2.1 mm, 1.7 mm particle size, 100 Å pore size, Phenomenex, Torrance, USA) was employed with a SecurityGuard ULTRA C18 cartridges (2.1 mm ID; 30 °C column temperature). The mobile phases (0.5 mL/min flow rate) were 0.1 % formic acid in both water (A) and acetonitrile (B) with the following gradient: 0-1 min 5 % B, 1-7 min 5-100 % B, 7-7.5 min 100 % B, 7.5-8 min 100-5% B, 8-10 min 5% B. Mass spectrometer was operated under the data dependent acquisition mode with positive heated electrospray ionization. The source parameters were: sheath gas flow, 53 AU; auxiliary gas flow, 14 AU; sweep gas flow, 3 AU; auxiliary gas temperature, 400 °C; spray voltage, 3.5 kV; inlet capillary temperature, 269 °C; S-lens level, 50 V. For data-dependent acquisition, MS1 scan was performed at m/z 100-1500 with the following parameters: resolution, 35,000 at m/z 200; maximum ion injection time, 100 ms; automated gain control (AGC) target, 5.0E5. For MS/MS acquisition, up to 5 MS/MS spectra per MS1 scan were recorded with the following parameters: resolution, 35,000 at m/z 200; maximum ion injection time, 100 ms; AGC target, 5.0E5; MS/MS precursor isolation window, m/z 3; isolation offset, m/z 0.5; normalized collision energy, a stepwise increase from 20 to 30 to 40 %; minimum AGC for MS/MS spectrum, 5.0E3; apex trigger, 2 to 15 s; dynamic precursor exclusion, 10 s. Obtained the raw data (.raw) was converted to .mzML format using MSConvert.

### Metabolite annotation

Raw data and mzML files are deposited in the MassIVE data repository (http://massive.ucsd.edu) under the accession number MSV000090088. Feature extraction was performed using MZmine 3.2.8^109^. The data was filtered by removing all MS/MS fragment ions within +/- 17 Da of the precursor m/z. MS/MS spectra were window filtered by choosing only the top 6 fragment ions in the +/- 50 Da window throughout the spectrum. A feature based molecular network (FBMN)^110^ was then created setting precursor ion mass tolerance to 0.02 Da and a MS/MS fragment ion tolerance of 0.02 Da. Edges were filtered to have a modified cosine score above 0.7 and more than 5 matched peaks. Further, edges between two nodes were kept in the network if and only if each of the nodes appeared in each other’s respective top 10 most similar nodes. Finally, the maximum size of a molecular family was set to 100, and the lowest scoring edges were removed from molecular families until the molecular family size was below this threshold.

The MZmine parameters were as follows - mass detection (parameters: MS1 noise level 3E5, MS2 noise level 0), ADAP chromatogram builder (parameters: minimum group size in number of scans 4, group intensity threshold 9E5, minimum highest intensity 3E6, 0.005 m/z tolerance or 10 ppm, retention time 0.88-7.5 min), local minimum resolver (parameters: chromatographic threshold 85%, minimum search range 0.05, minimum relative height 0.0%, minimum absolute height 1.0E3, min ratio of peak top/edge 1.7, peak duration range (min/mobility) 0-2.00, min number of data points 4), isotopic peaks grouper (m/z tolerance 0.01 m/z or 10 ppm, RT tolerance 0.30 min, maximum charge 5), join aligner (m/z tolerance 0.01 m/z or 10 ppm, weight for m/z 3, RT tolerance 0.10 min, weight for RT 1), gap filler (intensity tolerance 20%, m/z tolerance 0.001 m/z or 5.00 ppm, RT tolerance 0.1 min, minimum data points 2).

The spectra in the network were then searched against GNPS2’s spectral libraries^111^. The library spectra were filtered in the same manner as the input data. All matches kept between network spectra and library spectra were required to have a score above 0.7 and at least 6 matched peaks. The final annotation table produced 2754 features of which the GNPS had 413 level 2 annotations.

### Metabolite analysis

The annotations and hits per sample were analyzed in R 4.1.1 (R Foundation for Statistical Computing, Vienna, Austria). A cluster of samples (n=29) were found which were grown in a different batch of BHI media, and these samples were removed from subsequent analysis resulting in 84 *B. fragilis* isolates (commensal, n=38, extra-intestinal, n=31, unknown, n=18). The samples were normalized through the robust centered log ratio transformation (using the decostand function of the package vegan v2.6)^112^, which divides each value by the geometric mean of the observed features before applying the log transformation. Zeros are excluded from this transformation. The samples were corrected for the media blanks by subtracting the media blanks from the sample counts. We created a PCA plot of metabolites produced by the prcomp function of the normalized, corrected metabolite abundances. Violin plots were created using ggplot2 of media-corrected normalized metabolite production. The Wilcoxon rank-sum test was conducted to determine differential abundance of media-corrected and normalized metabolomics features between extra-intestinal and commensal strains. We determined significance with a Benjamini, Hochberg adjusted p-value ≤ 0.05 and log2-fold change ≥ 0.25. We calculated differential abundance of metabolites in all strains against the media controls with a student’s t-test with a significance cutoff of a Benjamini, Hochberg adjusted p-value ≤ 0.1 and log2-fold change ≥ 0.25. We created a heatmap in R with eucledian clustering of both rows (metabolites) and columns (isolates).

## Tables

**Table 1.** Newly sequenced Chu lab strains metadata.

**Table 2.** Accession identifiers for all public *Bacteroides fragilis* strains used in this study.

**Table 3.** Annotation for all unique genes in the *Bacteroides fragilis* pangenome.

**Table 4.** Phylogroup designation for each strain.

**Table 5.** Quality information on all isolates analyzed.

**Table 6.** Capsular polysaccharide and lipid compositions.

**Table 7.** Minimal inhibitory concentration (MIC, μg/mL) gradient of antibiotics tested against *B. fragilis* strains.

**Table 8.** Annotation of all genes identified through GWAS as significantly associated with isolation source.

**Table 9.** Primers for species-specific qPCR.

## Supplementary

**Supplemental Figure 1.**
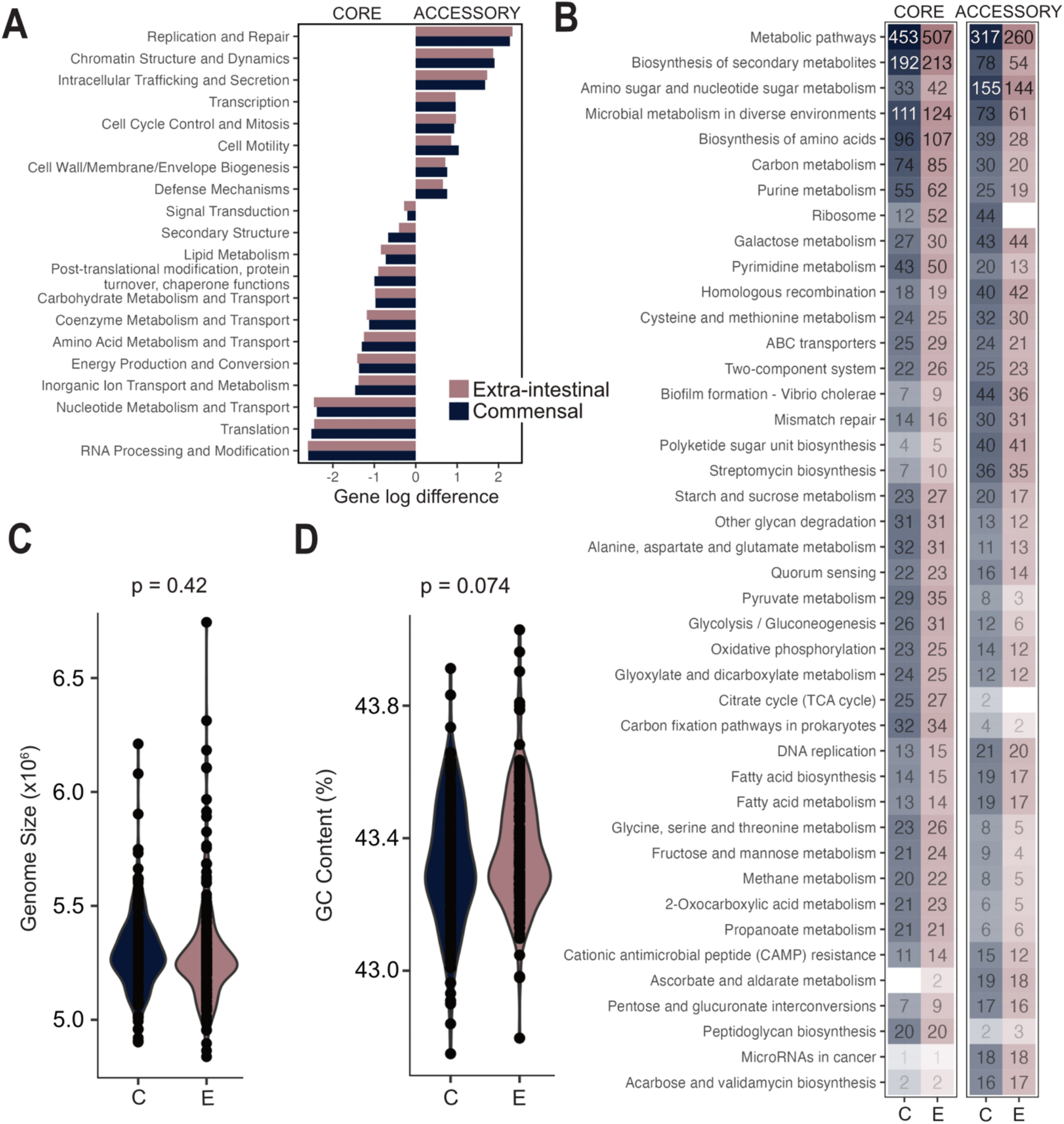
Genetic diversity among commensal and extra-intestinal *B. fragilis* isolates. **A)** Bar graph of log_2_ odds ratio between accessory and core genes per COG category separated by commensal (navy) versus extra-intestinal (pink) strains. **B)** The top 30 core and top 30 accessory KEGG categories, ordered by most frequently observed KEGG KO category. C, commensal; E, extra-intestinal. **C)** Genome size of commensal (C, n=281) and extra-intestinal (E, n=280) isolates, excluding assemblies derived from metagenomic assemblies, Welch’s t-test, p=0.42. **D)** GC content of commensal (C, n=281) and extra-intestinal (E, n=280) isolates, excluding assemblies derived from metagenomic assemblies, Welch’s t-test, p=0.074.

**Supplemental Figure 2.**
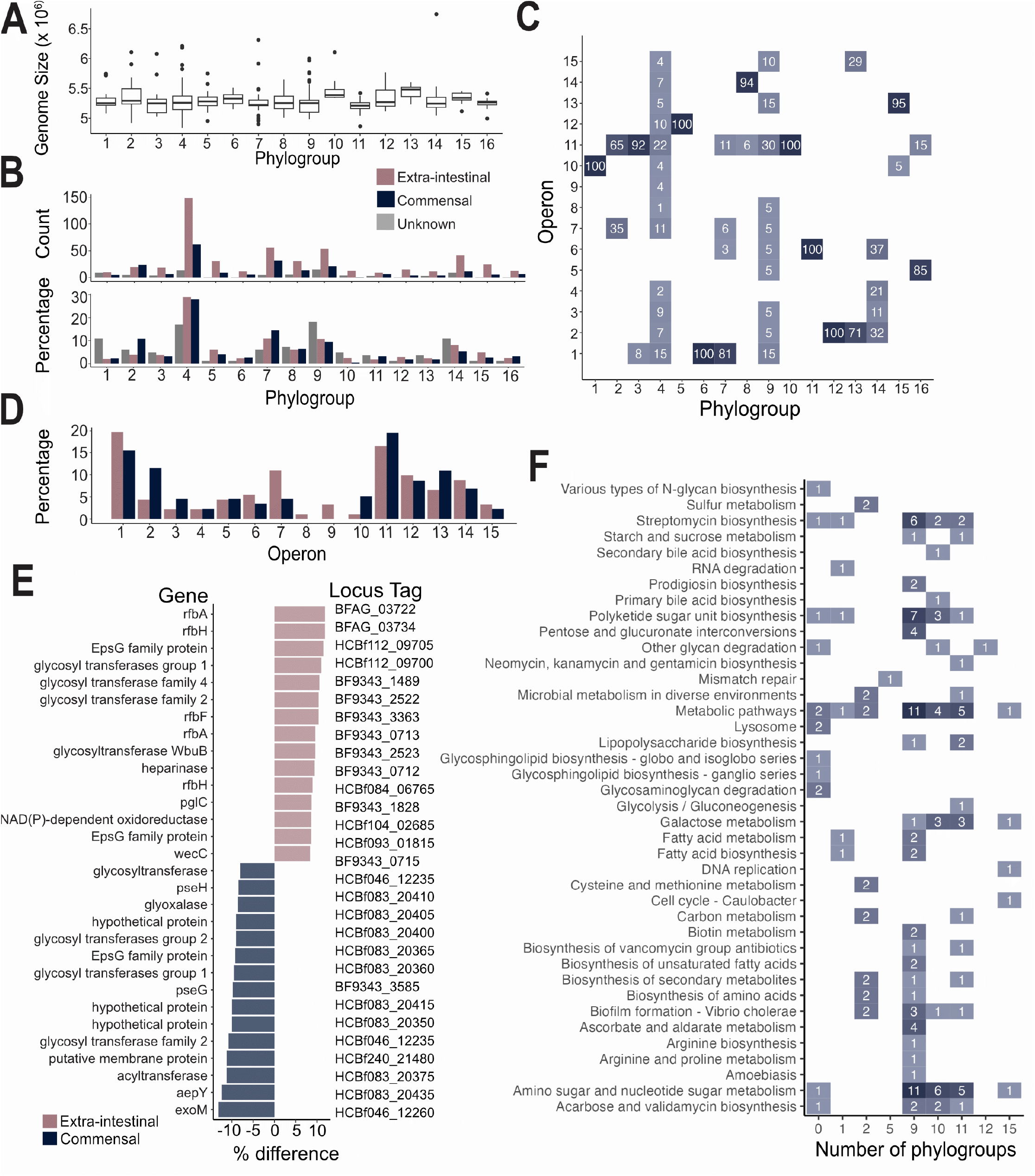
*B. fragilis* commensal and extra-intestinal strain distribution among phylogroups and niche specificity traits. **A)** Genome size of phylogroups 1-16. No statistically significant phylogroups by Welch’s t-test at a corrected p-value of 0.05. **B)** Distribution of *B. fragilis* isolates by isolation source across phylogroups. No statistically significant difference between isolation sources per phylogroup after chi-squared test with correction for multiple hypothesis testing. **C)** Distribution of phylogroups per PSA operon structure expressed in percentage. **D)** Distribution of commensal versus extra-intestinal *B. fragilis* isolates across PSA operons. **E)** Top 15 capsular polysaccharide genes which are differentially present in commensal compared with extra-intestinal strains. **F)** KEGG categories of genes core to a number of phylogroups (x-axis).

**Supplemental Figure 3.**
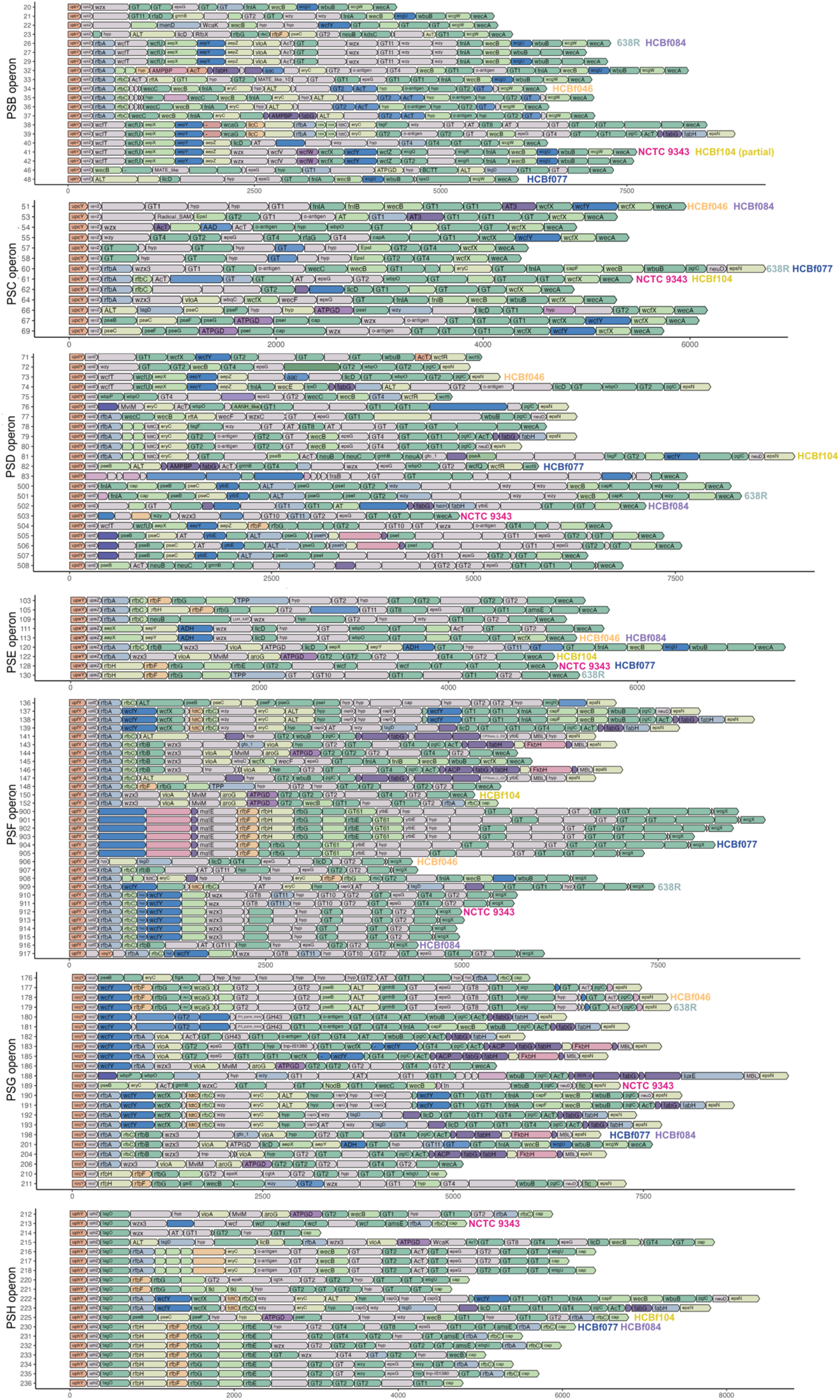
Polysaccharide operon structures among *B. fragilis* strains. PSB-H operon structure per all high-quality isolates in the *B. fragilis* pangenome (n=262). Genes are annotated with gene names, if annotated by bakta. Colored by COG category, annotated with information on select strains profiled throughout the study.

**Supplemental Figure 4.**
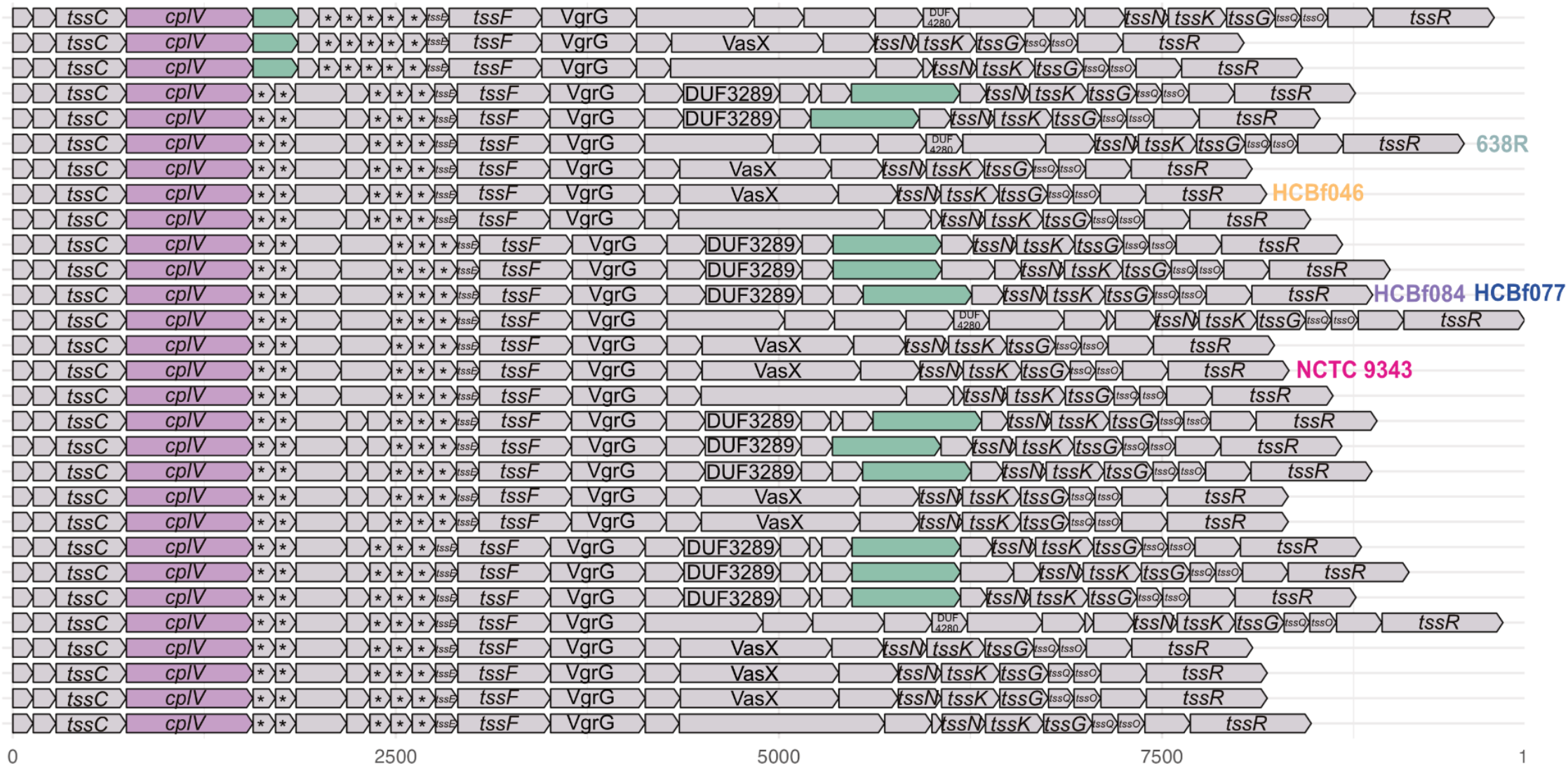
T6SSiii GA3 operon structures present in *B. fragilis* isolates. T6SSiii GA3 operon structure per all isolates in the *B. fragilis* pangenome (n=813), genes annotated with gene names if annotated by bakta, colored by COG category (grey, unknown; green, cell wall/membrane/envelope biogenesis; pink, replication and repair), annotated with information on phylogroup composition per operon.

**Supplemental Figure 5.**
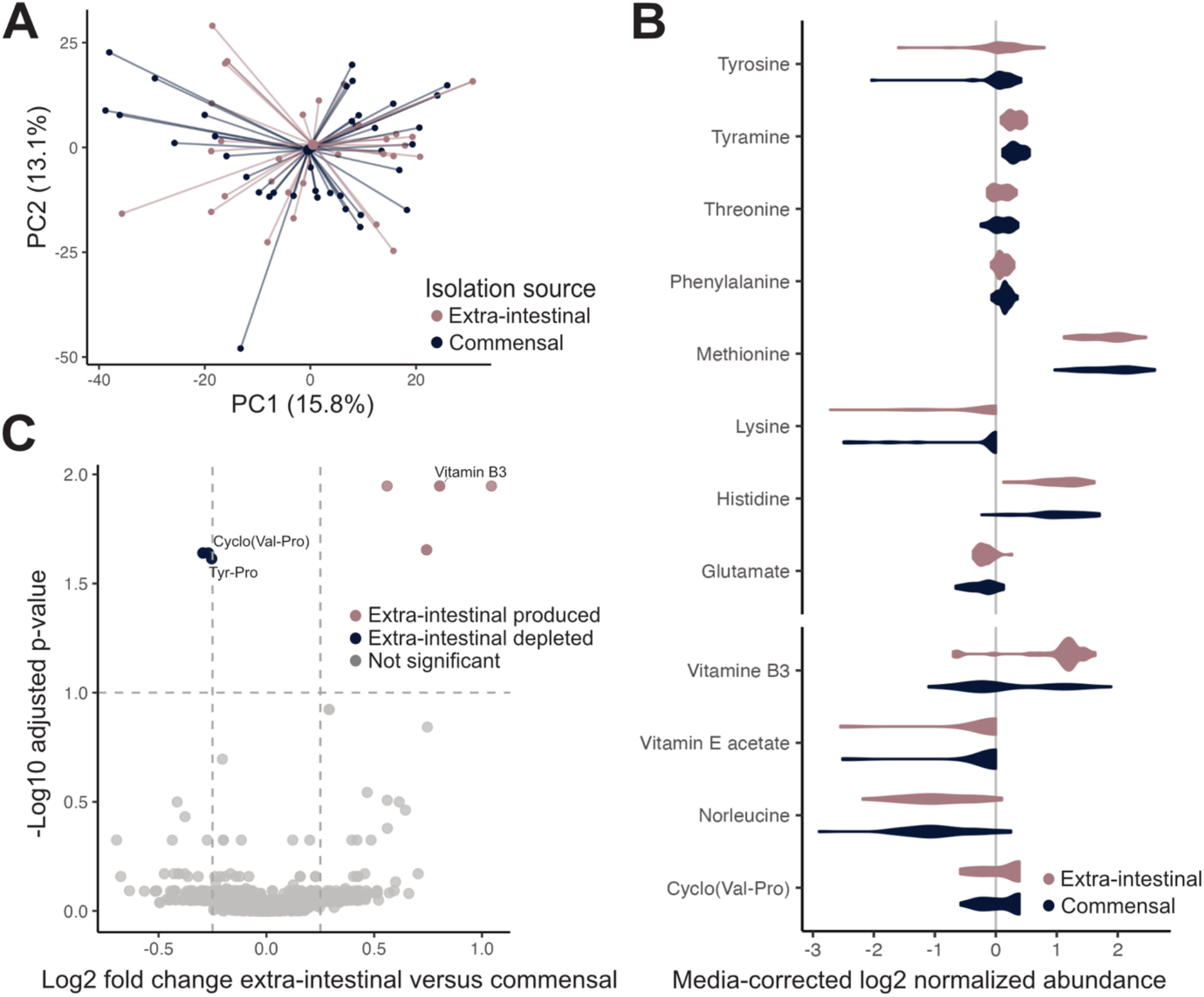
Metabolic features among commensal and extra-intestinal *B. fragilis*. **A)** PCA plot of media-corrected normalized abundances of strains (total, n=69; extra-intestinal=38; commensal=31) connected to the centroid of each group, colored by isolation source, (PERMANOVA, p=0.992). **B)** Metabolite production of annotated metabolites across *B. fragilis* strains normalized against the media control (n=73), comparing extra-intestinal (n=33) and commensal (n=40) strains. **C)** Volcano plot of differentially abundant metabolites of extra-intestinal (n=33) compared with commensal (n=40) strains. P-values on the y-axis are from a Wilcoxon rank-sum test comparing the average abundance of metabolites across strains versus a media control (dashed lines at Benjamini-Hochberg adjusted p-value ≤ 0.1 and log2-fold change ≥ 0.25).

**Supplemental Figure 6.**
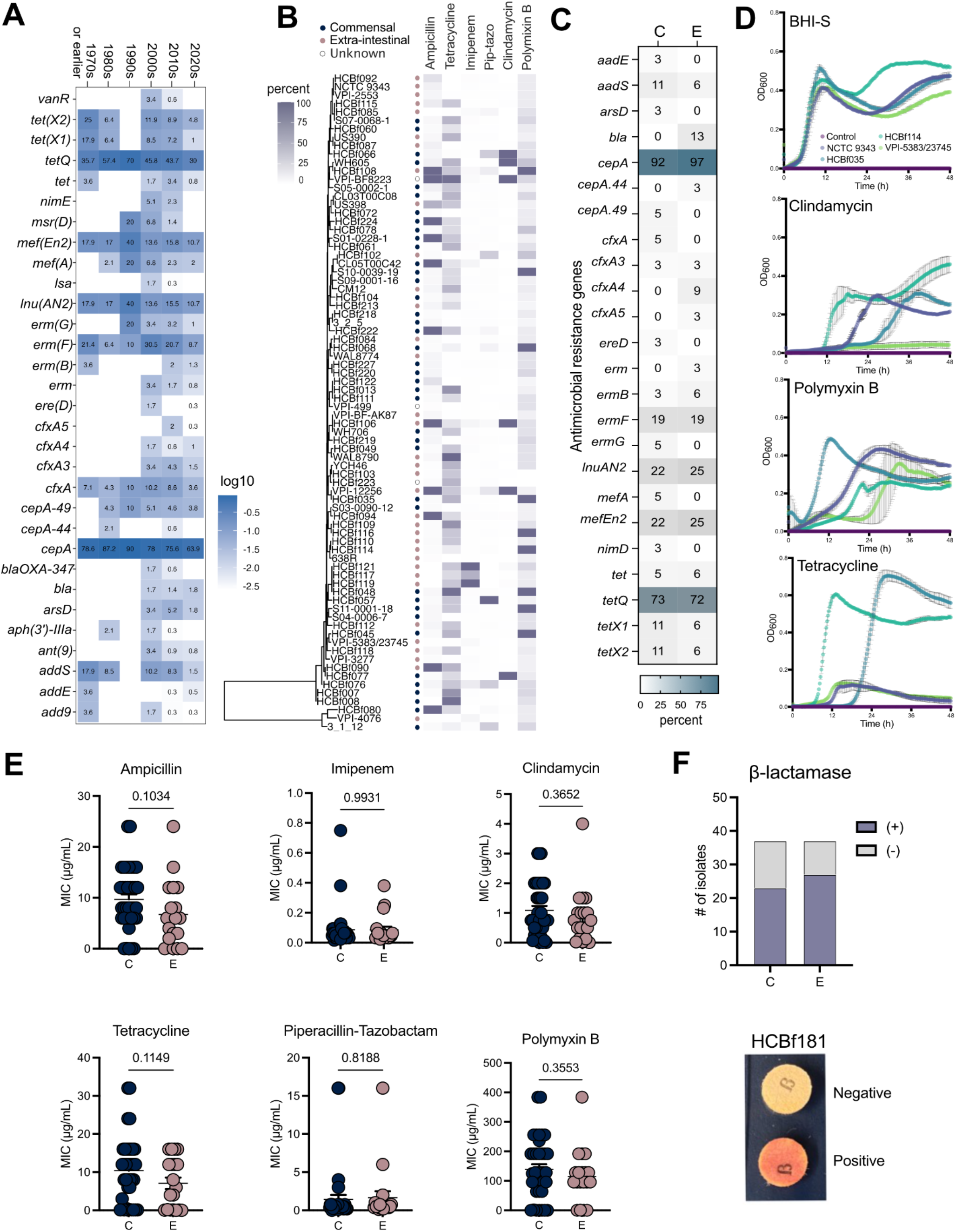
Antimicrobial resistance profile among commensal and pathogenic *B. fragilis*. **A)** The percentage of antimicrobial gene presence-absence, grouping *B. fragilis* isolates by decade of isolation (1970s or earlier, n=28; 1980s, n=47; 1990s, n=10; 2000s, n=348; 2010s, n=393; 2020s, n=393). **B)** Phylogenetic tree of *B. fragilis* strains used in antimicrobial resistance testing with proportion of resistance to ampicillin, tetracycline, imipenem, piperacillin-tazobactam (pip-tazo), clindamycin and polymyxin B. **C)** Percent of antimicrobial resistance gene distribution among *B. fragilis* strains (commensal, n=37; extra-intestinal, n=32). Unknowns were excluded from the analysis. **D)** Growth rate of representative strains in no antibiotic media control, clindamycin (0.03125 μg/mL), polymyxin B (400 μg/mL), and tetracycline (0.1875 μg/mL). **E)** Comparison of antimicrobial minimal inhibitory concentration (MIC, μg/mL) among commensal (C) and extra-intestinal (E) *B. fragilis* strains. Unknowns were excluded from the analysis. Unpaired t-test with Welch’s correction. **F)** Beta-lactamase production among commensal (C) and extra-intestinal (E) *B. fragilis* strains and representative positive reaction for beta-lactamase production (n=37 per group). p=0.457 from a Fisher’s exact test.

## Notes

### Competing Interest Statement

The authors have declared no competing interest.

## REFERENCES

1. Yassour, M., Vatanen, T., Siljander, H., Hämäläinen, A.-M., Härkönen, T., Ryhänen, S.J., Franzosa, E.A., Vlamakis, H., Huttenhower, C., Gevers, D., et al. (2016). Natural history of the infant gut microbiome and impact of antibiotic treatments on strain-level diversity and stability. Sci. Transl. Med. 8, 343ra81. 10.1126/scitranslmed.aad0917.

2. Mitchell, C.M., Mazzoni, C., Hogstrom, L., Bryant, A., Bergerat, A., Cher, A., Pochan, S., Herman, P., Carrigan, M., Sharp, K., et al. (2020). Delivery Mode Affects Stability of Early Infant Gut Microbiota. Cell Rep. Med. 1, 100156. 10.1016/j.xcrm.2020.100156.

3. Lee, S.M., Donaldson, G.P., Mikulski, Z., Boyajian, S., Ley, K., and Mazmanian, S.K. (2013). Bacterial colonization factors control specificity and stability of the gut microbiota. Nature 501, 426–429. 10.1038/nature12447.

4. Zhao, S., Lieberman, T.D., Poyet, M., Kauffman, K.M., Gibbons, S.M., Groussin, M., Xavier, R.J., and Alm, E.J. (2019). Adaptive evolution within the gut microbiomes of healthy people. Cell Host Microbe 25, 656–667.e8. 10.1016/j.chom.2019.03.007.

5. Kasper, D.L., Onderdonk, A.B., Polk, B.F., Bartlett, J.G. (1980). Surface Antigens as Virulence Factors in Infection with Bacteroides Fragilis. In Anaerobic Bacteria (Springer, Boston, MA).

6. Wexler, H.M. (2007). Bacteroides: the Good, the Bad, and the Nitty-Gritty. Clin. Microbiol. Rev. 20, 593– 621. 10.1128/CMR.00008-07.

7. Wallace, M.J., Jean, S., Wallace, M.A., Burnham, C.-A.D., and Dantas, G. (2022). Comparative Genomics of Bacteroides fragilis Group Isolates Reveals Species-Dependent Resistance Mechanisms and Validates Clinical Tools for Resistance Prediction. mBio 13, e03603–21. 10.1128/mbio.03603-21.

8. Carrow, H.C., Batachari, L.E., and Chu, H. (2020). Strain diversity in the microbiome: Lessons from Bacteroides fragilis. PLOS Pathog. 16, e1009056. 10.1371/journal.ppat.1009056.

9. Onderdonk, A.B., Kasper, D.L., Cisneros, R.L., and Bartlett, J.G. (1977). The Capsular Polysaccharide of Bacteroides fragilis as a Virulence Factor: Comparison of the Pathogenic Potential of Encapsulated and Unencapsulated Strains. J. Infect. Dis. 136, 82–89. 10.1093/infdis/136.1.82.

10. Van Doorn, J., Mooi, F.R., Verweij-van Vught, A.M.J.J., and MacLaren, D.M. (1987). Characterization of fimbriae from Bacteroides fragilis. Microb. Pathog. 3, 87–95. 10.1016/0882-4010(87)90067-2.

11. Brook, I., and Myhal, M.L. (1991). Adherence of Bacteroides fragilis group species. Infect. Immun. 59, 742– 744. 10.1128/iai.59.2.742-744.1991.

12. Ferreira, R., Alexandre, M.C.F., Antunes, E.N.F., Pinhao, A.T., Moraes, S.R., Ferreira, M.C.S., and Domingues, R.M.C.P. (1999). Expression of Bacteroides fragilis virulence markers in vitro. J. Med. Microbiol. 48, 999–1004. 10.1099/00222615-48-11-999.

13. Robertson, K.P., Smith, C.J., Gough, A.M., and Rocha, E.R. (2006). Characterization of *Bacteroides fragilis* Hemolysins and Regulation and Synergistic Interactions of HlyA and HlyB. Infect. Immun. 74, 2304–2316. 10.1128/IAI.74.4.2304-2316.2006.

14. Zakharzhevskaya, N.B., Vanyushkina, A.A., Altukhov, I.A., Shavarda, A.L., Butenko, I.O., Rakitina, D.V., Nikitina, A.S., Manolov, A.I., Egorova, A.N., Kulikov, E.E., et al. (2017). Outer membrane vesicles secreted by pathogenic and nonpathogenic Bacteroides fragilis represent different metabolic activities. Sci. Rep. 7, 5008. 10.1038/s41598-017-05264-6.

15. Oyston, P.C.F., and Handley, P.S. (1990). Surface structures, haemagglutination and cell surface hydrophobicity of Bacteroides fragilis strains. J. Gen. Microbiol. 136, 941–948. 10.1099/00221287-136-5-941.

16. Bjornson, A.B. (1990). Role of Humoral Factors in Host Resistance to the Bacteroides fragilis Group. Clin. Infect. Dis. 12, S161–S168. 10.1093/clinids/12.Supplement_2.S161.

17. Coyne, M.J., Tzianabos, A.O., Mallory, B.C., Carey, V.J., Kasper, D.L., and Comstock, L.E. (2001). Polysaccharide Biosynthesis Locus Required for Virulence of *Bacteroides fragilis*. Infect. Immun. 69, 4342– 4350. 10.1128/IAI.69.7.4342-4350.2001.

18. Round, J.L., and Mazmanian, S.K. (2010). Inducible Foxp3+ regulatory T-cell development by a commensal bacterium of the intestinal microbiota. Proc. Natl. Acad. Sci. U. S. A. 107, 12204–12209. 10.1073/pnas.0909122107.

19. Myers, L.L., Shoop, D.S., Stackhouse, L.L., Newman, F.S., Flaherty, R.J., Letson, G.W., and Sack, R.B. (1987). Isolation of Enterotoxigenic Bacteroides fragilis from Humans with Diarrheat. 25, 4.

20. Franco, A.A., Cheng, R.K., Goodman, A., and Sears, C.L. (2002). Modulation of *bft* expression by the *Bacteroides fragilis* pathogenicity island and its flanking region. Mol. Microbiol. 45, 1067–1077. 10.1046/j.1365-2958.2002.03077.x.

21. Pierce, J.V., and Bernstein, H.D. (2016). Genomic Diversity of Enterotoxigenic Strains of Bacteroides fragilis. PLOS ONE 11, e0158171. 10.1371/journal.pone.0158171.

22. Pasolli, E., Asnicar, F., Manara, S., Zolfo, M., Karcher, N., Armanini, F., Beghini, F., Manghi, P., Tett, A., Ghensi, P., et al. (2019). Extensive Unexplored Human Microbiome Diversity Revealed by Over 150,000 Genomes from Metagenomes Spanning Age, Geography, and Lifestyle. Cell 176, 649–662.e20. 10.1016/j.cell.2019.01.001.

23. Moraes, S.R., Gonçalves, R.B., Mouton, C., Seldin, L., Ferreira, M.C., and Domingues, R.M. (1999). Bacteroides fragilis isolates compared by AP-PCR. Res. Microbiol. 150, 257–263. 10.1016/s0923-2508(99)80050-3.

24. Salyers, A.A., Shoemaker, N.B., and Li, L.Y. (1995). In the driver’s seat: the Bacteroides conjugative transposons and the elements they mobilize. J. Bacteriol. 177, 5727–5731.

25. Shoemaker, N.B., Vlamakis, H., Hayes, K., and Salyers, A.A. (2001). Evidence for Extensive Resistance Gene Transfer among *Bacteroides* spp. and among *Bacteroides* and Other Genera in the Human Colon. Appl. Environ. Microbiol. 67, 561–568. 10.1128/AEM.67.2.561-568.2001.

26. Patrick, S. (2022). A tale of two habitats: Bacteroides fragilis, a lethal pathogen and resident in the human gastrointestinal microbiome. Microbiology 168. 10.1099/mic.0.001156.

27. Segerman, B. (2012). The genetic integrity of bacterial species: the core genome and the accessory genome, two different stories. Front. Cell. Infect. Microbiol. 2. 10.3389/fcimb.2012.00116.

28. Evanovich, E., Mattos, P.J. de S.M., and Guerreiro, J.F. (2022). Comprehensive Comparative Genomic revels: Bacteroides fragilis is a reservoir of antibiotic resistance genes in the gut microbiota. Preprint at bioRxiv, 10.1101/2022.05.30.494044.

29. Wozniak, R.A.F., and Waldor, M.K. (2010). Integrative and conjugative elements: mosaic mobile genetic elements enabling dynamic lateral gene flow. Nat. Rev. Microbiol. 8, 552–563. 10.1038/nrmicro2382.

30. Picard, B., Garcia, J.S., Gouriou, S., Duriez, P., Brahimi, N., Bingen, E., Elion, J., and Denamur, E. (1999). The link between phylogeny and virulence in Escherichia coli extraintestinal infection. Infect. Immun. 67, 546–553. 10.1128/IAI.67.2.546-553.1999.

31. De Filippis, F., Pasolli, E., Tett, A., Tarallo, S., Naccarati, A., De Angelis, M., Neviani, E., Cocolin, L., Gobbetti, M., Segata, N., et al. (2019). Distinct Genetic and Functional Traits of Human Intestinal Prevotella copri Strains Are Associated with Different Habitual Diets. Cell Host Microbe 25, 444–453.e3. 10.1016/j.chom.2019.01.004.

32. White, H., Vos, M., Sheppard, S.K., Pascoe, B., and Raymond, B. (2022). Signatures of selection in core and accessory genomes indicate different ecological drivers of diversification among Bacillus cereus clades. Mol. Ecol. 31, 3584–3597. 10.1111/mec.16490.

33. Lagerstrom, K.M., and Hadly, E.A. (2023). Under-Appreciated Phylogroup Diversity of *Escherichia coli* within and between Animals at the Urban-Wildland Interface. Appl. Environ. Microbiol. 89, e00142–23. 10.1128/aem.00142-23.

34. Shen, Y., Torchia, M.L.G., Lawson, G.W., Karp, C.L., Ashwell, J.D., and Mazmanian, S.K. (2012). Outer Membrane Vesicles of a Human Commensal Mediate Immune Regulation and Disease Protection. Cell Host Microbe 12, 509–520. 10.1016/j.chom.2012.08.004.

35. Chu, H., Khosravi, A., Kusumawardhani, I.P., Kwon, A.H.K., Vasconcelos, A.C., Cunha, L.D., Mayer, A.E., Shen, Y., Wu, W.-L., Kambal, A., et al. (2016). Gene-Microbiota Interactions Contribute to the Pathogenesis of Inflammatory Bowel Disease. Science 352, 1116–1120. 10.1126/science.aad9948.

36. Gibson, F.C., III, Onderdonk, A.B., Kasper, D.L., and Tzianabos, A.O. (1998). Cellular Mechanism of Intraabdominal Abscess Formation by Bacteroides fragilis1. J. Immunol. 160, 5000–5006. 10.4049/jimmunol.160.10.5000.

37. Wang, Y., Kalka-Moll, W.M., Roehrl, M.H., and Kasper, D.L. (2000). Structural basis of the abscess-modulating polysaccharide A2 from *Bacteroides fragilis*. Proc. Natl. Acad. Sci. 97, 13478–13483. 10.1073/pnas.97.25.13478.

38. Neff, C.P., Rhodes, M.E., Arnolds, K.L., Collins, C.B., Donnelly, J., Nusbacher, N., Jedlicka, P., Schneider, J.M., McCarter, M.D., Shaffer, M., et al. (2016). Diverse Intestinal Bacteria Contain Putative Zwitterionic Capsular Polysaccharides with Anti-inflammatory Properties. Cell Host Microbe 20, 535–547. 10.1016/j.chom.2016.09.002.

39. Husain, F., Tang, K., Veeranagouda, Y., Boente, R., Patrick, S., Blakely, G., and Wexler, H.M. (2017). Novel large-scale chromosomal transfer in Bacteroides fragilis contributes to its pan-genome and rapid environmental adaptation. Microb. Genomics 3, e000136. 10.1099/mgen.0.000136.

40. Lindberg, A.A., Weintraub, A., and Zihringer, U. (1990). Structure-Activity Relationships in Lipopolysaccharides of Bacteroides fragiiis. Rev Infect Dis. 10.1093/clinids/12.supplement_2.s133.

41. Jacobson, A.N., Choudhury, B.P., and Fischbach, M.A. (2018). The Biosynthesis of Lipooligosaccharide from *Bacteroides thetaiotaomicron*. mBio 9, e02289–17. 10.1128/mBio.02289-17.

42. Reeves, P. (1995). Role of O-antigen variation in the immune response. Trends in Microbiology 3, 381– 386. 10.1016/s0966-842x(00)88983-0.

43. Coyne, M.J., and Comstock, L.E. (2019). Type VI secretion systems and the gut microbiota. Microbiol. Spectr. 7, 10.1128/microbiolspec.PSIB-0009–2018.

44. Zhang, Z.J., Cole, C.G., Coyne, M.J., Lin, H., Dylla, N., Smith, R.C., Waligurski, E., Ramaswamy, R., Woodson, C., Burgo, V., et al. (2024). Comprehensive analyses of a large human gut Bacteroidales culture collection reveal species and strain level diversity and evolution. Preprint, 10.1101/2024.03.08.584156.

45. Casterline, B.W., Hecht, A.L., Choi, V.M., and Bubeck Wardenburg, J. (2017). The *Bacteroides fragilis* pathogenicity island links virulence and strain competition. Gut Microbes 8, 374–383. 10.1080/19490976.2017.1290758.

46. Hill, C.A. (2024). Bacteroides fragilis toxin expression enables lamina propria niche acquisition in the developing mouse gut. Nat. Microbiol. 9.

47. Coyne, M.J., Roelofs, K.G., and Comstock, L.E. (2016). Type VI secretion systems of human gut Bacteroidales segregate into three genetic architectures, two of which are contained on mobile genetic elements. BMC Genomics 17, 58. 10.1186/s12864-016-2377-z.

48. Wexler, A.G., Bao, Y., Whitney, J.C., Bobay, L.-M., Xavier, J.B., Schofield, W.B., Barry, N.A., Russell, A.B., Tran, B.Q., Goo, Y.A., et al. (2016). Human symbionts inject and neutralize antibacterial toxins to persist in the gut. Proc. Natl. Acad. Sci. 113, 3639–3644. 10.1073/pnas.1525637113.

49. Verster, A.J., Ross, B.D., Radey, M.C., Bao, Y., Goodman, A.L., Mougous, J.D., and Borenstein, E. (2017). The Landscape of Type VI Secretion across Human Gut Microbiomes Reveals Its Role in Community Composition. Cell Host Microbe 22, 411–419.e4. 10.1016/j.chom.2017.08.010.

50. Chatzidaki-Livanis, M., Geva-Zatorsky, N., and Comstock, L.E. (2016). Bacteroides fragilis type VI secretion systems use novel effector and immunity proteins to antagonize human gut Bacteroidales species. Proc. Natl. Acad. Sci. U. S. A. 113, 3627–3632. 10.1073/pnas.1522510113.

51. Jiang, K., Li, W., Tong, M., Xu, J., Chen, Z., Yang, Y., Zang, Y., Jiao, X., Liu, C., Lim, B., et al. (2023). Bacteroides fragilis ubiquitin homologue drives intraspecies bacterial competition in the gut microbiome. Nat. Microbiol. 9, 70–84. 10.1038/s41564-023-01541-5.

52. Pudlo, N.A., Urs, K., Crawford, R., Pirani, A., Atherly, T., Jimenez, R., Terrapon, N., Henrissat, B., Peterson, D., Ziemer, C., et al. (2022). Phenotypic and Genomic Diversification in Complex Carbohydrate-Degrading Human Gut Bacteria. mSystems 7, e0094721. 10.1128/msystems.00947-21.

53. Buckwold, S.L., Shoemaker, N.B., Sears, C.L., and Franco, A.A. (2007). Identification and Characterization of Conjugative Transposons CTn86 and CTn9343 in *Bacteroides fragilis* Strains. Appl. Environ. Microbiol. 73, 53–63. 10.1128/AEM.01669-06.

54. Franco, A.A. (2004). The *Bacteroides fragilis* Pathogenicity Island Is Contained in a Putative Novel Conjugative Transposon. J. Bacteriol. 186, 6077–6092. 10.1128/JB.186.18.6077-6092.2004.

55. Prindiville, T.P., Sheikh, R.A., Cohen, S.H., Tang, Y.J., Cantrell, M.C., and Silva, J. (2000). Bacteroides fragilis enterotoxin gene sequences in patients with inflammatory bowel disease. Emerg. Infect. Dis. 6, 171– 174. 10.3201/eid0602.000210.

56. Toprak, N.U., Yagci, A., Gulluoglu, B.M., Akin, M.L., Demirkalem, P., Celenk, T., and Soyletir, G. (2006). A possible role of Bacteroides fragilis enterotoxin in the aetiology of colorectal cancer. Clin. Microbiol. Infect. Off. Publ. Eur. Soc. Clin. Microbiol. Infect. Dis. 12, 782–786. 10.1111/j.1469-0691.2006.01494.x.

57. Sears, C.L. (2009). Enterotoxigenic Bacteroides fragilis: a Rogue among Symbiotes. Clin. Microbiol. Rev. 22, 349–369. 10.1128/cmr.00053-08.

58. McMillan, A.S., Foley, M.H., Perkins, C.E., and Theriot, C.M. (2024). Loss of *Bacteroides thetaiotaomicron* bile acid-altering enzymes impacts bacterial fitness and the global metabolic transcriptome. Microbiol. Spectr. 12, e03576–23. 10.1128/spectrum.03576-23.

59. Rimal, B., Collins, S.L., Tanes, C.E., Rocha, E.R., Granda, M.A., Solanki, S., Hoque, N.J., Gentry, E.C., Koo, I., Reilly, E.R., et al. (2024). Bile salt hydrolase catalyses formation of amine-conjugated bile acids. Nature 626, 859–863. 10.1038/s41586-023-06990-w.

60. García-Bayona, L., Coyne, M.J., and Comstock, L.E. (2021). Mobile Type VI secretion system loci of the gut Bacteroidales display extensive intra-ecosystem transfer, multi-species spread and geographical clustering. PLOS Genet. 17, e1009541. 10.1371/journal.pgen.1009541.

61. Sheahan, M.L., Coyne, M.J., Flores, K., Garcia-Bayona, L., Chatzidaki-Livanis, M., Sundararajan, A., Holst, A.Q., Barquera, B., and Comstock, L.E. (2023). A ubiquitous mobile genetic element disarms a bacterial antagonist of the gut microbiota. Preprint at bioRxiv, 10.1101/2023.08.25.553775.

62. Wexler, H.M., Tenorio, E., and Pumbwe, L. (2009). Characteristics of Bacteroides fragilis lacking the major outer membrane protein, OmpA. Microbiol. Read. Engl. 155, 2694–2706. 10.1099/mic.0.025858-0.

63. Sydenham, T.V., Overballe-Petersen, S., Hasman, H., Wexler, H., Kemp, M., and Justesen, U.S. (2019). Complete hybrid genome assembly of clinical multidrug-resistant Bacteroides fragilis isolates enables comprehensive identification of antimicrobial-resistance genes and plasmids. Microb. Genomics 5, e000312. 10.1099/mgen.0.000312.

64. Wright, D.P., Rosendale, D.I., and Roberton, A.M. (2000). *Prevotella* enzymes involved in mucin oligosaccharide degradation and evidence for a small operon of genes expressed during growth on mucin. FEMS Microbiol. Lett. 190, 73–79. 10.1111/j.1574-6968.2000.tb09265.x.

65. Csiszovszki, Z., Krishna, S., Orosz, L., Adhya, S., and Semsey, S. (2011). Structure and Function of the D-Galactose Network in Enterobacteria. mBio 2, e00053–11. 10.1128/mBio.00053-11.

66. Treangen, T.J., and Rocha, E.P.C. (2011). Horizontal Transfer, Not Duplication, Drives the Expansion of Protein Families in Prokaryotes. PLOS Genet. 7, e1001284. 10.1371/journal.pgen.1001284.

67. Salyers, A., Gupta, A., and Wang, Y. (2004). Human intestinal bacteria as reservoirs for antibiotic resistance genes. Trends Microbiol. 12, 412–416. 10.1016/j.tim.2004.07.004.

68. Kurokawa, K., Itoh, T., Kuwahara, T., Oshima, K., Toh, H., Toyoda, A., Takami, H., Morita, H., Sharma, V.K., Srivastava, T.P., et al. (2007). Comparative Metagenomics Revealed Commonly Enriched Gene Sets in Human Gut Microbiomes. DNA Res. 14, 169–181. 10.1093/dnares/dsm018.

69. Groussin, M., Poyet, M., Sistiaga, A., Kearney, S.M., Moniz, K., Noel, M., Hooker, J., Gibbons, S.M., Segurel, L., Froment, A., et al. (2021). Elevated rates of horizontal gene transfer in the industrialized human microbiome. Cell 184, 2053–2067.e18. 10.1016/j.cell.2021.02.052.

70. Blandford, L.E., Johnston, E.L., Sanderson, J.D., Wade, W.G., and Lax, A.J. (2019). Promoter orientation of the immunomodulatory *Bacteroides fragilis* capsular polysaccharide A (PSA) is off in individuals with inflammatory bowel disease (IBD). Gut Microbes 10, 569–577. 10.1080/19490976.2018.1560755.

71. Russell, A.B., Wexler, A.G., Harding, B.N., Whitney, J.C., Bohn, A.J., Goo, Y.A., Tran, B.Q., Barry, N.A., Zheng, H., Peterson, S.B., et al. (2014). A Type VI Secretion-Related Pathway in Bacteroidetes Mediates Interbacterial Antagonism. Cell Host Microbe 16, 227–236. 10.1016/j.chom.2014.07.007.

72. Hill, C. (2012). Virulence or Niche Factors: What’s in a Name? J. Bacteriol. 194, 5725–5727. 10.1128/JB.00980-12.

73. Franco, A.A., Cheng, R.K., Chung, G.-T., Wu, S., Oh, H.-B., and Sears, C.L. (1999). Molecular Evolution of the Pathogenicity Island of Enterotoxigenic *Bacteroides fragilis* Strains. J. Bacteriol. 181, 6623–6633. 10.1128/JB.181.21.6623-6633.1999.

74. Rasko, D.A., Rosovitz, M.J., Myers, G.S.A., Mongodin, E.F., Fricke, W.F., Gajer, P., Crabtree, J., Sebaihia, M., Thomson, N.R., Chaudhuri, R., et al. (2008). The Pangenome Structure of *Escherichia coli* : Comparative Genomic Analysis of *E. coli* Commensal and Pathogenic Isolates. J. Bacteriol. 190, 6881– 6893. 10.1128/JB.00619-08.

75. Terhes, G., Brazier, J.S., Sóki, J., Urbán, E., and Nagy, E. (2007). Coincidence of bft and cfiA genes in a multi-resistant clinical isolate of Bacteroides fragilis. J. Med. Microbiol. 56, 1416–1418. 10.1099/jmm.0.47242-0.

76. Sóki, J., Eitel, Z., Terhes, G., Nagy, E., and Urbán, E. (2013). Occurrence and analysis of rare cfiA–bft doubly positive Bacteroides fragilis strains. Anaerobe 23, 70–73. 10.1016/j.anaerobe.2013.06.008.

77. Fogarty, E.C., Schechter, M.S., Lolans, K., Sheahan, M.L., Veseli, I., Moore, R.M., Kiefl, E., Moody, T., Rice, P.A., Yu, M.K., et al. (2024). A cryptic plasmid is among the most numerous genetic elements in the human gut. Cell 187, 1206–1222.e16. 10.1016/j.cell.2024.01.039.

78. Smith, C.J. (1985). Development and use of cloning systems for Bacteroides fragilis: cloning of a plasmid-encoded clindamycin resistance determinant. J. Bacteriol. 164, 294–301. 10.1128/jb.164.1.294-301.1985.

79. J.L. Johson and D.A. Ault (1978). Taxonomy of the Bacteroides II. Correlation of Phenotypic Characteristics with Deoxyribonucleic Acid Homology Groupings for Bacteroides fragilis and Other Saccharolytic Bacteroides Species. INT. J. SYST. BACTERIO 28, 257–268. https://doi.org/0020-7713/78/0028-0257$02.00/0.

80. Frank, J.A., Reich, C.I., Sharma, S., Weisbaum, J.S., Wilson, B.A., and Olsen, G.J. (2008). Critical Evaluation of Two Primers Commonly Used for Amplification of Bacterial 16S rRNA Genes. Appl. Environ. Microbiol. 74, 2461–2470. 10.1128/AEM.02272-07.

81. Shaffer, J.P., Marotz, C, Belda-Ferre, P., Martino, C, Wandro, S, Estaki, M., Salido, R.A., Carpenter, C.S, Zaramela, L.S, Minich, J.J., et al. (2021). A comparison of DNA/RNA extraction protocols for high-throughput sequencing of microbial communities. Biotechniques 70, 149–159. 10.2144/btn-2020-0153.

82. Sanders, J.G., Nurk, S., Salido, R.A., Minich, J., Xu, Z.Z., Zhu, Q., Martino, C., Fedarko, M., Arthur, T.D., Chen, F., et al. (2019). Optimizing sequencing protocols for leaderboard metagenomics by combining long and short reads. Genome Biol. 20, 226. 10.1186/s13059-019-1834-9.

83. Brennan, C., Salido, R.A., Belda-Ferre, P., Bryant, M., Cowart, C., Tiu, M.D., González, A., McDonald, D., Tribelhorn, C., Zarrinpar, A., et al. (2023). Maximizing the potential of high-throughput next-generation sequencing through precise normalization based on read count distribution. mSystems 8, e0000623. 10.1128/msystems.00006-23.

84. Chen, S., Zhou, Y., Chen, Y., and Gu, J. (2018). fastp: an ultra-fast all-in-one FASTQ preprocessor. Bioinformatics 34, i884–i890. 10.1093/bioinformatics/bty560.

85. Li, H. (2018). Minimap2: pairwise alignment for nucleotide sequences. Bioinformatics 34, 3094–3100. 10.1093/bioinformatics/bty191.

86. Gonzalez, A., Navas-Molina, J.A., Kosciolek, T., McDonald, D., Vázquez-Baeza, Y., Ackermann, G., DeReus, J., Janssen, S., Swafford, A.D., Orchanian, S.B., et al. (2018). Qiita: rapid, web-enabled microbiome meta-analysis. Nat. Methods 15, 796–798. 10.1038/s41592-018-0141-9.

87. Renee Oles (2023). rolesucsd/Panpiper. (GitHub repository).

88. Seemann, T. (2022). Shovill.

89. Schwengers, O., Jelonek, L., Dieckmann, M.A., Beyvers, S., Blom, J., and Goesmann, A. (2021). Bakta: rapid and standardized annotation of bacterial genomes via alignment-free sequence identification. Microb. Genomics 7, 000685. 10.1099/mgen.0.000685.

90. Feldgarden, M., Brover, V., Gonzalez-Escalona, N., Frye, J.G., Haendiges, J., Haft, D.H., Hoffmann, M., Pettengill, J.B., Prasad, A.B., Tillman, G.E., et al. (2021). AMRFinderPlus and the Reference Gene Catalog facilitate examination of the genomic links among antimicrobial resistance, stress response, and virulence. Sci. Rep. 11, 12728. 10.1038/s41598-021-91456-0.

91. Cantalapiedra, C.P., Hernández-Plaza, A., Letunic, I., Bork, P., and Huerta-Cepas, J. (2021). eggNOG-mapper v2: Functional Annotation, Orthology Assignments, and Domain Prediction at the Metagenomic Scale. Mol. Biol. Evol. 38, 5825–5829. 10.1093/molbev/msab293.

92. Tonkin-Hill, G., MacAlasdair, N., Ruis, C., Weimann, A., Horesh, G., Lees, J.A., Gladstone, R.A., Lo, S., Beaudoin, C., Floto, R.A., et al. (2020). Producing polished prokaryotic pangenomes with the Panaroo pipeline. Genome Biol. 21, 180. 10.1186/s13059-020-02090-4.

93. Tonkin-Hill, G., Gladstone, R.A., Pöntinen, A.K., Arredondo-Alonso, S., Bentley, S.D., and Corander, J. (2023). Robust analysis of prokaryotic pangenome gene gain and loss rates with Panstripe. Genome Res. 33, 129–140. 10.1101/gr.277340.122.

94. Price, M.N., Dehal, P.S., and Arkin, A.P. (2010). FastTree 2 – Approximately Maximum-Likelihood Trees for Large Alignments. PLOS ONE 5, e9490. 10.1371/journal.pone.0009490.

95. Minh, B.Q., Schmidt, H.A., Chernomor, O., Schrempf, D., Woodhams, M.D., von Haeseler, A., and Lanfear, R. (2020). IQ-TREE 2: New Models and Efficient Methods for Phylogenetic Inference in the Genomic Era. Mol. Biol. Evol. 37, 1530–1534. 10.1093/molbev/msaa015.

96. Stamatakis, A. (2014). RAxML version 8: a tool for phylogenetic analysis and post-analysis of large phylogenies. Bioinformatics 30, 1312–1313. 10.1093/bioinformatics/btu033.

97. Ondov, B.D., Treangen, T.J., Melsted, P., Mallonee, A.B., Bergman, N.H., Koren, S., and Phillippy, A.M. (2016). Mash: fast genome and metagenome distance estimation using MinHash. Genome Biol. 17, 132. 10.1186/s13059-016-0997-x.

98. Abram, K., Udaondo, Z., Bleker, C., Wanchai, V., Wassenaar, T.M., Robeson, M.S., and Ussery, D.W. (2021). Mash-based analyses of Escherichia coli genomes reveal 14 distinct phylogroups. Commun. Biol. 4, 1–12. 10.1038/s42003-020-01626-5.

99. Lees, J.A., Galardini, M., Bentley, S.D., Weiser, J.N., and Corander, J. (2018). pyseer: a comprehensive tool for microbial pangenome-wide association studies. Bioinforma. Oxf. Engl. 34, 4310–4312. 10.1093/bioinformatics/bty539.

100. Shannon, P., Markiel, A., Ozier, O., Baliga, N.S., Wang, J.T., Ramage, D., Amin, N., Schwikowski, B., and Ideker, T. (2003). Cytoscape: A Software Environment for Integrated Models of Biomolecular Interaction Networks. Genome Res. 13, 2498–2504. 10.1101/gr.1239303.

101. Westphal O and Jann K (1965). Bacterial lipopolysaccharides extraction with phenolwater and further applications of the procedure. Methods in Carbohydrate Chemistry *Vol.* 5, 92–93.

102. Martens, E.C., Chiang, H.C., and Gordon, J.I. (2008). Mucosal Glycan Foraging Enhances Fitness and Transmission of a Saccharolytic Human Gut Bacterial Symbiont. Cell Host Microbe 4, 447–457. 10.1016/j.chom.2008.09.007.

103. Blazanin, M. (2023). gcplyr: an R package for microbial growth curve data analysis. Preprint, 10.1101/2023.04.30.538883.

104. Sprouffske, K., and Wagner, A. (2016). Growthcurver: an R package for obtaining interpretable metrics from microbial growth curves. BMC Bioinformatics 17, 172. 10.1186/s12859-016-1016-7.

105. RStudio Team (2020). RStudio: Integrated Development Environment for R. Version 2024.04.0+735 (RStudio, PBC).

106. Neal, M., Thiruppathy, D., and Zengler, K. (2023). Genome-scale metabolic modeling of the human gut bacterium Bacteroides fragilis strain 638R. PLOS Comput. Biol. 19, e1011594. 10.1371/journal.pcbi.1011594.

107. King, Z.A., Lu, J., Dräger, A., Miller, P., Federowicz, S., Lerman, J.A., Ebrahim, A., Palsson, B.O., and Lewis, N.E. (2016). BiGG Models: A platform for integrating, standardizing and sharing genome-scale models. Nucleic Acids Res. 44, D515–D522. 10.1093/nar/gkv1049.

108. Heirendt, L., Arreckx, S., Pfau, T., Mendoza, S.N., Richelle, A., Heinken, A., Haraldsdóttir, H.S., Wachowiak, J., Keating, S.M., Vlasov, V., et al. (2019). Creation and analysis of biochemical constraint-based models using the COBRA Toolbox v.3.0. Nat. Protoc. 14, 639–702. 10.1038/s41596-018-0098-2.

109. Schmid, R., Heuckeroth, S., Korf, A., Smirnov, A., Myers, O., Dyrlund, T.S., Bushuiev, R., Murray, K.J., Hoffmann, N., Lu, M., et al. (2023). Integrative analysis of multimodal mass spectrometry data in MZmine 3. Nat. Biotechnol. 41, 447–449. 10.1038/s41587-023-01690-2.

110. Nothias, L.-F., Petras, D., Schmid, R., Dührkop, K., Rainer, J., Sarvepalli, A., Protsyuk, I., Ernst, M., Tsugawa, H., Fleischauer, M., et al. (2020). Feature-based molecular networking in the GNPS analysis environment. Nat. Methods 17, 905–908. 10.1038/s41592-020-0933-6.

111. Wang, M., Carver, J.J., Phelan, V.V., Sanchez, L.M., Garg, N., Peng, Y., Nguyen, D.D., Watrous, J., Kapono, C.A., Luzzatto-Knaan, T., et al. (2016). Sharing and community curation of mass spectrometry data with Global Natural Products Social Molecular Networking. Nat. Biotechnol. 34, 828–837. 10.1038/nbt.3597.

112. Martino, C., Morton, J.T., Marotz, C.A., Thompson, L.R., Tripathi, A., Knight, R., and Zengler, K. (2019). A Novel Sparse Compositional Technique Reveals Microbial Perturbations. mSystems 4, e00016–19. 10.1128/mSystems.00016-19.

